# A multi-omics genome-and-transcriptome single-cell atlas of human preimplantation embryogenesis reveals the cellular and molecular impact of chromosome instability

**DOI:** 10.1101/2023.03.08.530586

**Authors:** Elia Fernandez Gallardo, Alejandro Sifrim, Joel Chappell, Jonas Demeulemeester, Jennifer Clara Herrmann, Robin Vermotte, Alison Kerremans, Michiel Van der Haegen, Jens Van Herck, Sebastiaan Vanuytven, Katy Vandereyken, Iain C. Macaulay, Joris Robert Vermeesch, Karen Peeraer, Sophie Debrock, Vincent Pasque, Thierry Voet

**Author notes:** Authors contributed equally (Elia Fernandez Gallardo, Alejandro Sifrim). Corresponding, co-last authors (V.P.), (T.V.).

## Abstract

The frequent acquisition of genomic abnormalities in human preimplantation embryos is a leading cause of pregnancy loss, but does not necessarily prohibit healthy offspring. However, the impact of genomic abnormalities on cellular states and development of the early human embryo remains largely unclear. Here, we characterise aneuploidy and reconstruct gene regulatory networks in human preimplantation embryos, and investigate gene expression and developmental perturbations instigated by aneuploidy using single-cell genome-and-transcriptome sequencing (G&T-seq). At the genomic level, we show that acquired numerical and structural chromosomal aberrations are frequent across all stages of early embryogenesis and in all cell lineages. At the transcriptome level, we identify regulators of cell identity and uncover a network of 248 transcription factors from 10 major gene regulatory modules that characterise the distinct lineages of human preimplantation embryos. By integrating single-cell DNA-with RNA-information, we unveil how expression levels are affected by losses or gains of the corresponding genes in embryonic cells across human preimplantation development, as well as how copy-number aberrant transcription factor genes perturb the expression of their cognate target genes in euploid regions. Furthermore, we reveal a majority of aneuploid cells show a developmental delay and reduced fitness, indicating cell competition within the mosaic diploid-aneuploid embryo, which may contribute to selection against aneuploid cells and the birth of healthy offspring from mosaic diploid-aneuploid embryos. In summary, our multi-modal analyses provide unprecedented insights into early human embryo development.

## INTRODUCTION

Preimplantation embryogenesis comprises the first cell cycles of life, an important 7-day period during which the totipotent fertilised egg gives rise to the first differentiated cell types for uterine implantation and development of an individual. Understanding this stage is paramount for discerning causes of infertility, implantation failure and spontaneous abortion, as well as developmental disorders. Although counter-intuitive, human preimplantation embryos are highly prone to chromosomal instability (CIN), not only following *in vitro* fertilisation (IVF)^1–5^, but also after natural conception^6–10^. Chromosomal gains and losses may arise during meiosis, affecting the whole embryo after fertilisation^1, 2, 6–12^. Even more frequently, numerical and structural chromosomal abnormalities can be acquired post-fertilisation during mitosis, leading to chromosomal mosaicism, in which genetically different cell lineages coexist in the same embryo^2–4^. However, the impact of acquired aneuploidies –which can be whole chromosome or segmental in nature– on gene expression programmes, cellular functional states and human embryo development remains largely unclear.

Acquired chromosomal and/or segmental aneuploidy –referred as (segmental) aneuploidy hereafter– can not only cause implantation failure and spontaneous abortion^8–10, 13^, but also genomic disorders due to mosaic or constitutional genomic aberrations^14–17^. Yet, mosaic (segmental) aneuploidies in the embryo do not always prohibit the development of normal offspring^4, 13, 18–20^. In an effort to maximise the success of IVF, many clinics have adopted the use of preimplantation genetic testing for aneuploidy (PGT-A)^21, 22^, whereby genetic analyses of a multicellular trophectoderm biopsy is used in an attempt to select euploid embryos with maximal developmental potential for uterine transfer^23^. However, since mosaicism for (segmental) aneuploidies can be compatible with healthy offspring, interpreting and counselling a mosaic PGT-A result remains highly challenging in the clinic. Different practices are applied across the world^24^ without a fundamental understanding of (segmental) aneuploid cell biology in the human embryo.

From studies in mammalian cell lines and yeast ^25–28^ mainly on bulk cell populations, we know that constitutional aneuploidies –i.e., present in all cells– may elicit gene dosage effects. These are changes in gene expression, both in *cis* and in *trans*, that result from changes in DNA copy-number of one or more genes. Gene dosage effects may have a major impact on development. They may result in cell stress response(s) characterised by reduced expression of genes involved in proliferation, nucleic-acid metabolism and protein processing, and by increased expression of genes involved in the endoplasmic reticulum, lysosomal pathways and membrane functions^25–28^. In mouse embryos with induced chromosomal instability and mosaicism for aneuploidy, it has been shown that aneuploid cells in the inner cell mass undergo apoptosis^29^, while in the trophectoderm aneuploid cells persist at lower cell cycle rate^30^. In addition, in human gastruloids, cell selection mechanisms that deplete aneuploid cells have been reported^31^. In human embryos with constitutional aneuploidy, different phenotypic outcomes can be present at blastocyst stage^32^ or post-implantation stages^33^ depending on the cell type^32^ and the nature of the aneuploidy^33^. However, despite their importance, the molecular and individual cellular responses to aneuploidies in human embryos remain largely unknown.

Recent advances in low-input and single-cell technologies provide an opportunity to study the changes in gene expression in the often mosaic human embryos, comprised of both (segmental) aneuploid and euploid cells. One study analysed human embryos up to the morula stage by partitioning embryo cells for genomic or other cells for gene expression analysis, and reported a gene-expression signature of aneuploidy-containing embryos before embryonic genome activation (EGA) at the 4-to 8-cell stage^34^. Other studies that inferred aneuploidies from single-cell RNA-seq data after EGA^35^ or that analysed low-input multicellular biopsies of human embryos, led to conflicting aneuploidy-induced gene expression signatures^32, 36–38^. Hence, the precise effects of (segmental) aneuploidies on gene expression dosage in *cis* and *trans*, and in turn on cellular responses, remain unknown in the transcriptionally dynamic and yet-to-be-determined gene regulatory landscape of human preimplantation embryogenesis. Consequently, it also remains unclear to what extent (segmental) aneuploidies may be inferred from single-cell transcriptomes across the entire human preimplantation period, necessitating a genome-and-transcriptome analysis at single-cell resolution.

Here, we apply genome-and-transcriptome sequencing (G&T-seq) on the same single cells of human preimplantation embryos to study the cell biology of (segmental) aneuploidy during early embryogenesis. We discover that autosomal DNA gains and losses result in gene dosage effects in *cis*, which differ across various stages of preimplantation embryo development and are not fully compensated. In addition, we provide a comprehensive analysis of the gene regulatory network operative during preimplantation human embryogenesis, and demonstrate how DNA copy number aberrations of autosomal transcription factor genes lead to the perturbation of their target gene expression in *trans*. Lastly, we detect cellular stress response(s) to aneuploidies concomitant with a developmental delay of aneuploid cells, and reveal cell competition to be operative, which may limit the contribution of aneuploid cells to mosaic embryos.

## RESULTS

### Single-cell G&T-seq enables profiling chromosome instability and gene expression in the same cells of human preimplantation embryos

To investigate the DNA and RNA of entire embryos at single-cell resolution across preimplantation development, we analysed 756 single cells from 112 human embryos spanning the first week after fertilisation using scG&T-seq^39^. In total, 370 genomes and 756 transcriptomes were sequenced, of which 295 and 576 were retained after stringent quality control (Methods), corresponding to 62 and 109 embryos, respectively (**Fig. 1a**). For 254 cells of 58 embryos across embryonic days E1-E6 (**Fig. 1b**) both high-quality DNA-and RNA-seq data were obtained (**Extended Data Table 1**).

**Figure 1.**
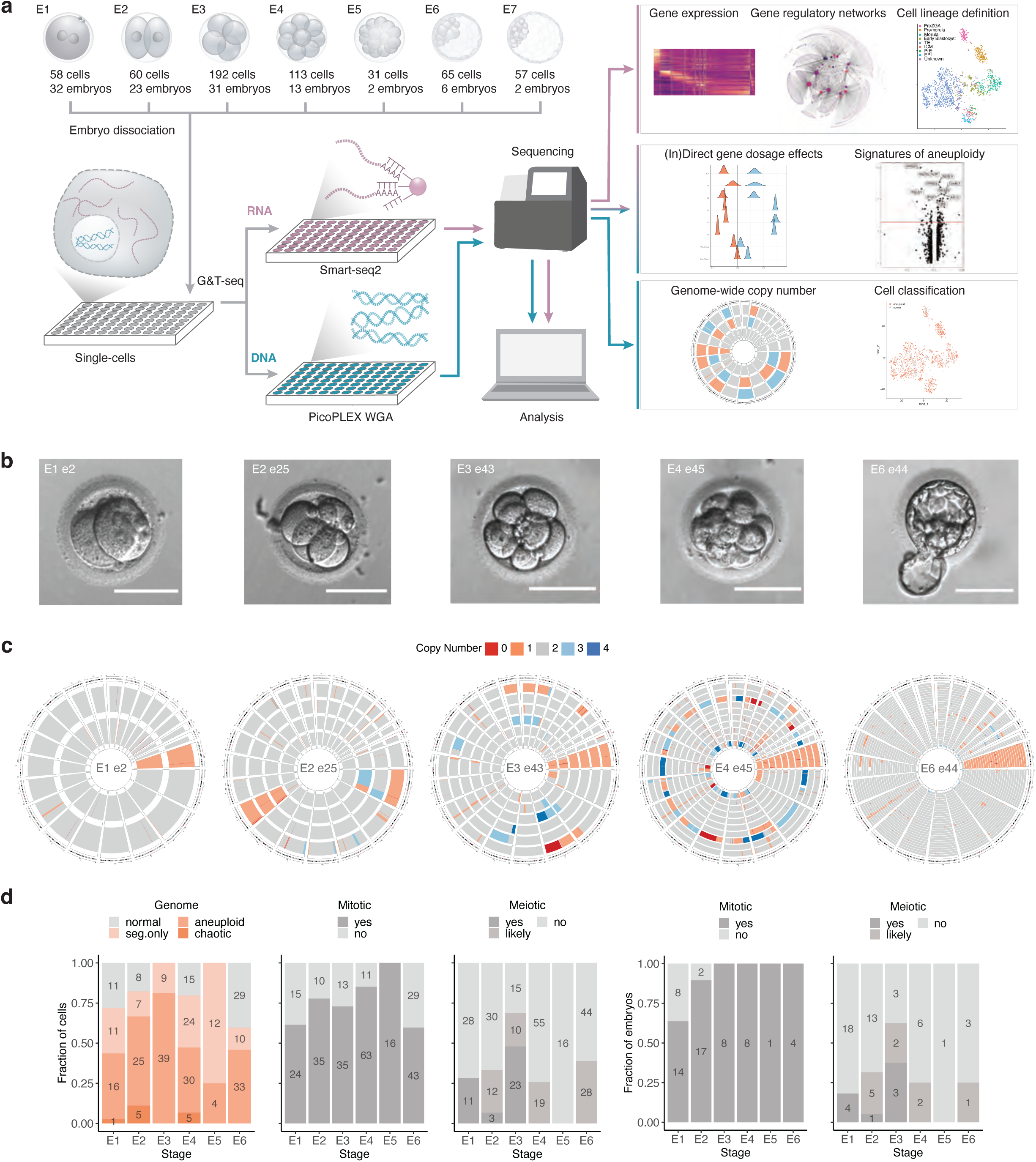
Characterisation of genomic instability during preimplantation development by single-cell G&T-seq. **a.** Experimental design, G&T-seq methodology and analysis, including also the number of embryos and cells with good quality data for each developmental stage, are depicted. **b.** Time lapse microscopy images of selected embryos prior to dissociation. Embryonic day of development and embryo ID is indicated on top. Scale bar corresponds to 100μm. **c.** Single-cell genome-wide copy number (CN) profiles of embryos in **b**. For each Circos plot, concentric circles correspond to individual cells in the embryo above. Each sector corresponds to a chromosome ordered clockwise from 1 to X. E=embryonic day, e=embryo identification number. **d.** Proportion and number of cells and embryos with different type and origin of genomic abnormalities.

Using DNA copy number analysis, genomes were classified into four categories: 1) normal, 2) aneuploid (i.e., one or more chromosomes are lost or gained and may be accompanied by additional segmental aberration(s)), 3) segmental aneuploid (i.e. only segment(s) of chromosome(s) are lost or gained), and 4) chaotic, (including segmental copy number abnormalities in more than half of the autosomal chromosomes) (Methods, **Fig. 1c, Extended Data Fig. 1a-b**). Of all cells, 50% were aneuploid, 25% had only segmental aneuploidies, and 4% presented a chaotic copy number landscape, affecting respectively, 63%, 39% and 10% of the embryos (**Fig. 1d, Extended Data Fig. 1c, Extended Data Table 2**). Putative meiotic abnormalities were observed in 18% of the embryos (36% of cells), while 92% of embryos (74% of cells) showed mitotic abnormalities. Normal cells (21%) were present in 39% of embryos and 15% of embryos were normal in all analysed cells (**Fig. 1d, Extended Data Fig. 1c, Extended Data Table 2**). These observations are consistent with previous single-cell genome surveys of human embryos^1, 35, 40–44^.

Next, we investigated the scG&T-derived transcriptomes of 576 cells (**Extended Data Fig. 2a**). Following dimensionality reduction and marker gene analyses, single-cell gene expression profiles exhibited a developmental trajectory from E1 to E7 (**Extended Data Fig. 2b**) and marker genes typical for pre-EGA, EGA and post-EGA embryonic stages (**Extended Data Fig. 2c,d**). We integrated the 576 single-cell transcriptomes with publicly available single-cell RNA-seq data from early human embryos^45^, totalling 2105 cells (**Extended Data Fig. 3a**). Our multi-omic scG&T-derived transcriptomes aligned well with mono-omic SmartSeq2-derived single-cell RNA-seq data and formed clusters that again reflected developmental progression and expressed known marker genes (**Fig. 2a, Extended Data Fig. 3a-c**). In summary, our single-cell G&T-seq data recapitulates both known genomic and transcriptomic characteristics of human preimplantation embryo development.

**Figure 2.**
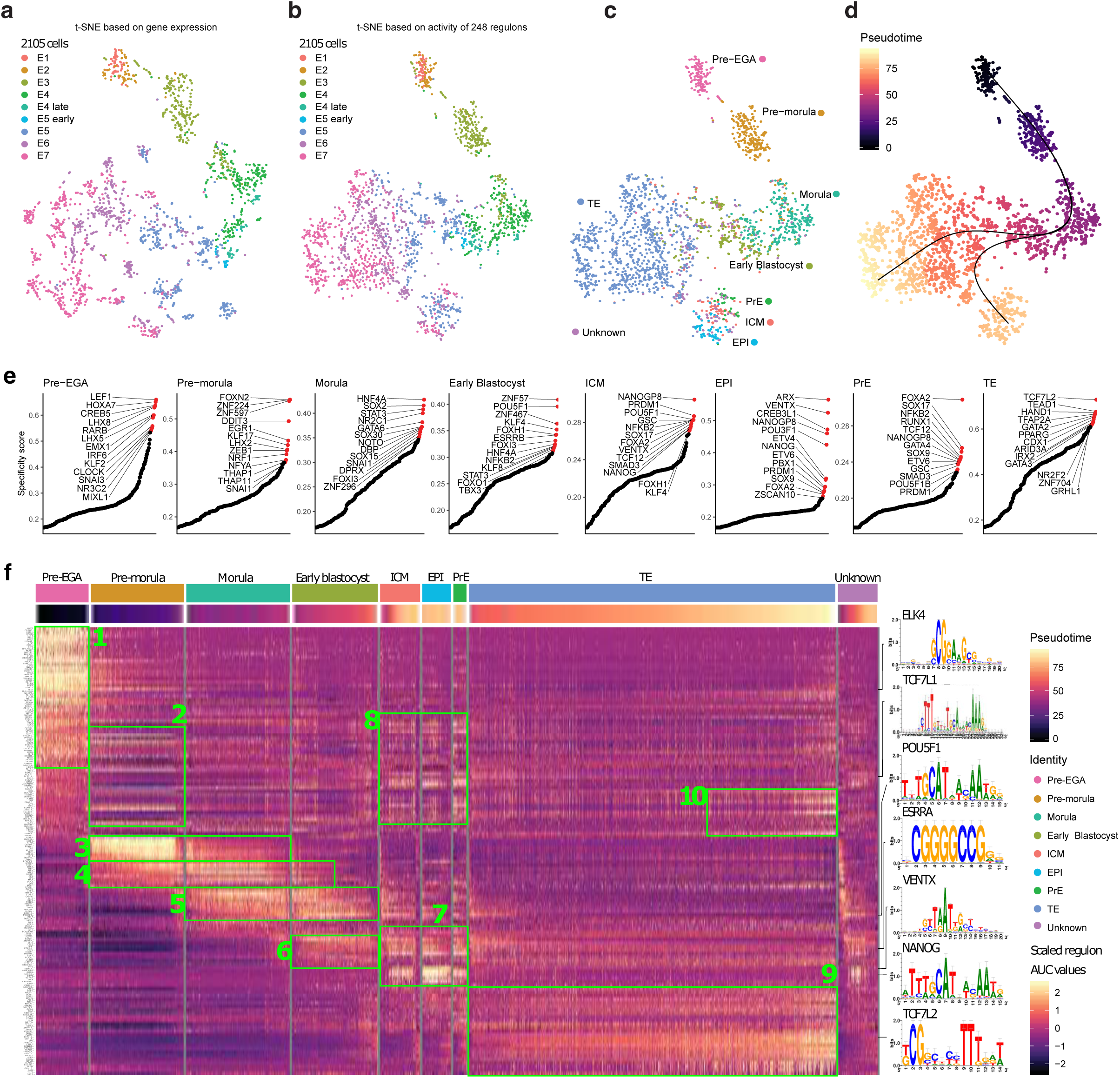
Gene regulatory networks active in human preimplantation embryos. **a.** t-SNE generated from integrated gene counts from 2105 single cells, embryonic day projected on top. **b-d.** t-SNE generated from regulon AUC values from 2105 single cells. Embryonic day, lineage and pseudotime projected on top, respectively. **e.** Regulon specificity ranking within each lineage. A high specificity score indicates that a regulon is highly specific to that lineage. **f.** Heat map displaying regulon expression (scaled regulon AUC values). The heat map is ordered by cell lineage and within each lineage ordered by pseudotime. Selected motifs for transcription factors are displayed on the right of the heat map. Green boxes indicate selected modules which group regulons expressed at similar time points or within a lineage.

**Figure 3.**
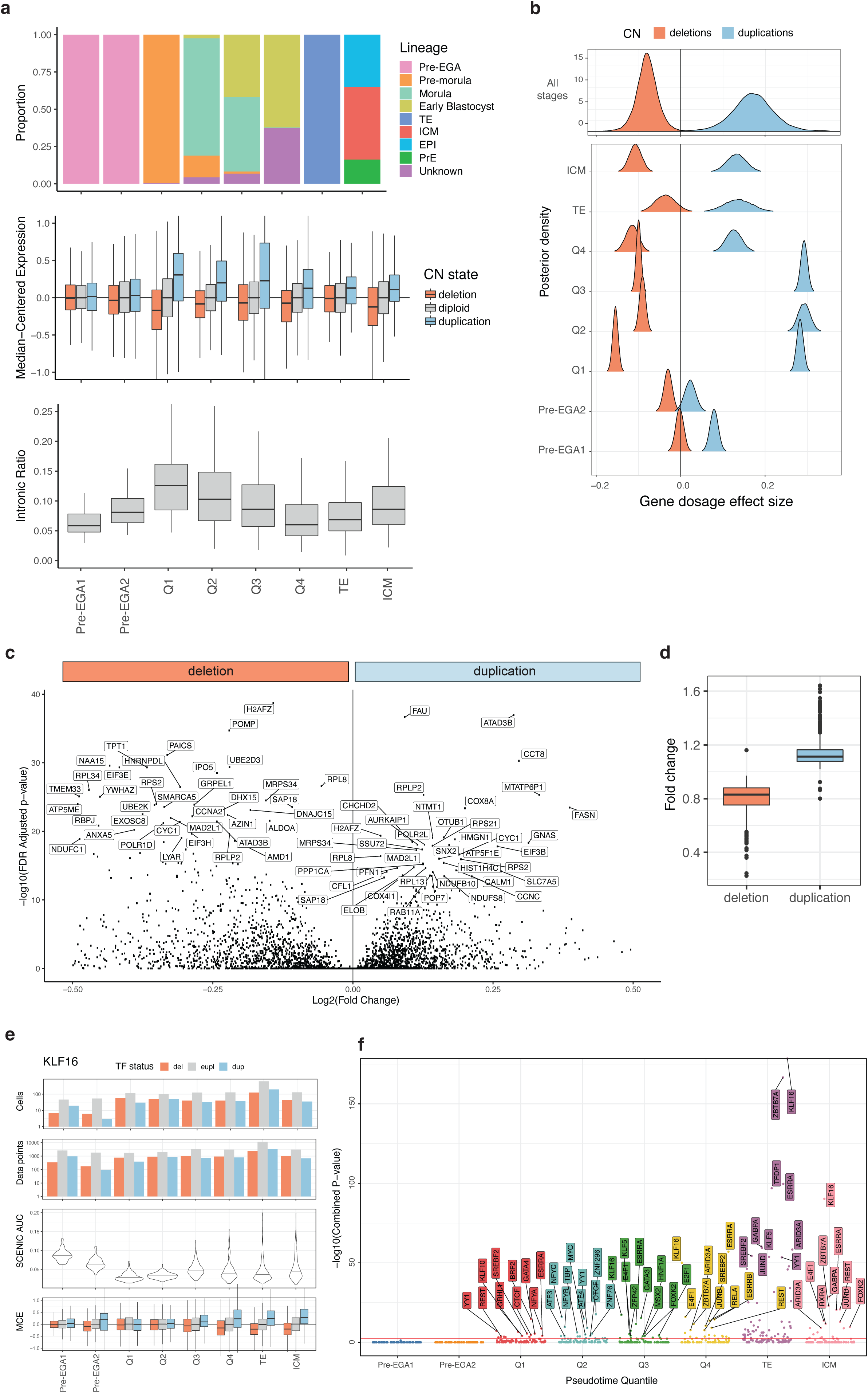
Dynamics of gene-wise dosage effects along human preimplantation development. **a.** Groups of transcriptionally similar cells defined by pseudotime quantiles and coloured by embryonic cell lineage (top). Direct gene dosage effects visualised by euploid normalised median-centred expression of genes in a duplication, deletion and normal copy number for cells in each pseudotime quantile (middle). Intronic read ratio of each pseudotime quantile indicative of active transcription levels (bottom). **b.** Gene dosage effects visualised by Bayesian posterior density distributions of the observed deviations from the median centred expression for genes in a deletion or duplication for each pseudotime quantile. **c.** Top 40 genes showing significant dosage effects during preimplantation embryo development. Their functions are described in the text and in **Extended Data Table 6. d.** Fold change distribution of dosage sensitive genes located in deletions and duplications. **e.** Indirect gene dosage effects of the transcription factor (TF) KLF16 on its target genes. For each pseudotime quantile and TF copy number, number of cells, number of euploid target genes (i.e., data points), regulon expression (i.e., SCENIC AUC) and normalised median-centred expression (MCE) of KLF16 target genes are indicated. **f.** For each pseudotime quantile, TFs showing significant indirect dosage effects on their target genes are depicted. Red line indicates the threshold p-value for a Wilcoxon ranked-sum test.

We expected gene expression to be strongly perturbed in aneuploid cells. However, euploid and aneuploid cells did not cluster separately (**Extended Data Fig. 2b**). These results suggest that aneuploidy does not induce dominant global changes in gene expression in all cells and that instead, changes in the transcriptome of preimplantation human embryos are mainly driven by developmental programs in both euploid and aneuploid cells.

### Reconstructing gene regulatory networks across human preimplantation development

To further scrutinise the potential functional impact of (segmental) aneuploidy in human embryos, we first reconstructed the gene regulatory networks of human preimplantation development. Gene regulatory networks (GRNs) govern cellular identity and fate transitions. Using pySCENIC, a method that links co-expression of genes and transcription factors (TFs) with cis-regulatory motif analysis to infer GRN activity in each cell^46, 47^, we identified 248 regulons, each consisting of a TF with its predicted downstream target genes. A *t*-SNE constructed from regulon activity grouped cells into distinct regulatory states that characterise cell states and fate transitions across the first week of human embryo development (**Fig. 2b-d, Extended Data Fig. 3d,e**). In agreement with previous studies^45, 48–54^, we detected high regulon activity for TFs in distinct cell types including maternally inherited LHX8 in pre-EGA cells, NFYA in the pre-morula, GATA6 in morula, POU5F1 in early blastocyst and inner cell mass (ICM), GATA2 in the trophectoderm (TE), NANOG in epiblast cells (EPI) and SOX17 in the primitive endoderm (PrE) (**Fig. 2e, Extended Data Fig. 3f**). Additionally, multiple regulons identified as active here, such as GATA6, GATA2, NANOG, POU5F1 and SOX2, were previously found to have their binding motif enriched in accessible chromatin in human preimplantation embryos^51, 55, 56^. Pseudotime analysis identified two branching trajectories corresponding to the TE in one branch, versus ICM, EPI and PrE lineages in the other branch, in line with the first cell fate decision segregating TE from the ICM in human embryos^45, 53, 54, 57–59^ (**Fig. 2d**). Altogether, we have comprehensively reconstructed the GRNs active throughout human preimplantation development (**Extended Data Fig. 4a**).

**Figure 4.**
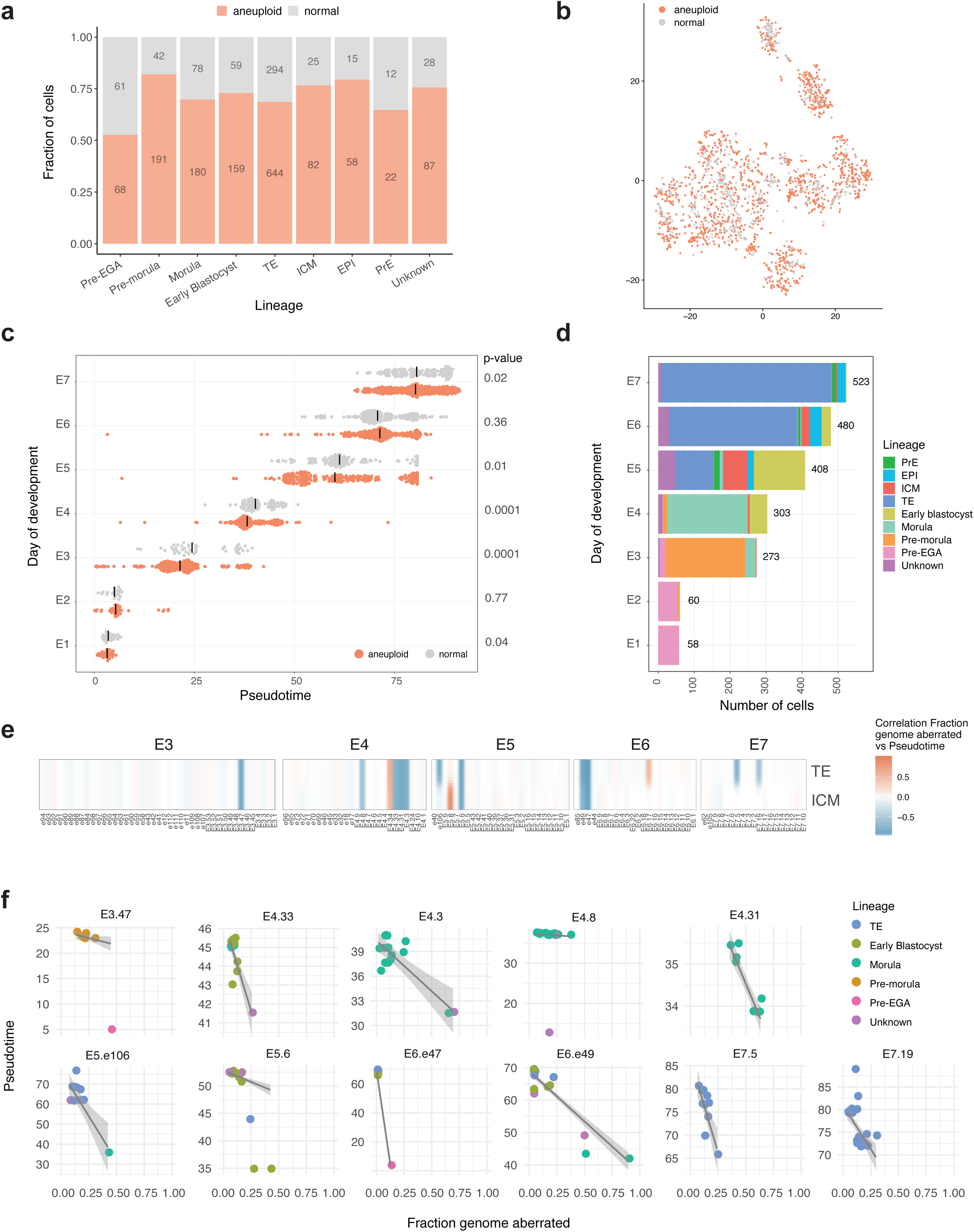
Developmental delay of aneuploid cells. **a.** Proportion and number of aneuploid cells per lineage. No preferential allocation of aneuploid cells was observed. A cell was classified as aneuploid if >60% of a chromosome arm was called as CN≠2 based on inferCNV or DNA-seq where available (methods). **b.** Regulon based t-SNE coloured by cell genomic status. **c.** Pseudotime values of normal and aneuploid cells on each day of development. Horizontal bars indicate the median pseudotime for the group. Significant p-values (<0.05) are indicative of developmental delay of aneuploid cells. **d.** Number of cells per day of embryonic development coloured by lineage. **e.** Heatmap showing the correlation value between the genomic fraction aberrated and pseudotime of cells in each embryo. Pseudotime splits the trajectory for ICM and TE lineages. **f.** Embryos which cells show a significant negative correlation between an abnormal fraction of the genome and pseudotime TE branch.

To predict which TFs may drive cell identity in human preimplantation embryogenesis, we assigned a specificity score for each regulon in each lineage, ranking regulons from most to least specific^60^ (**Fig. 2e**). We identified five classes of TFs: 1) TFs that have been demonstrated to be crucial in human preimplantation embryos such as POU5F1 in the early blastocyst^61, 62^, GATA3 in the trophectoderm^53^ and TEAD4 in the morula ^63^ (**Fig. 2f**), 2) TFs that are expected to be active based on immunofluorescence studies in human embryos or mouse studies such as e.g. GATA2 and NR2F2 in the trophectoderm^53, 54, 64, 65^, NANOG in the naive epiblast^48, 50, 54^ and SOX17 and FOXA2 in the primitive endoderm^49,54,65–67,^ 3) species-specific TFs such as VENTX, active in the human naive epiblast but absent in the mouse genome^57^, 4) TFs with species-specific temporal gene regulatory activity such as SOX2, active in the human morula but later in mouse embryos^49, 53, 68^, and 5) a large number of candidate TFs with unknown function in human embryos, which we propose might drive cell identity in human preimplantation embryos. These include ESRRB in the early blastocyst, POU3F1 in the epiblast and ARID3A in the trophectoderm (**Fig. 2e**). Several of these findings are not expected. For example, the regulon ARX, associated with forebrain development and myogenesis in the mouse^69, 70^, had the highest specificity score within the naïve epiblast. Similarly, the regulon HNF4A is the highest ranked regulon for the morula, while in the mouse this TF has been implicated in the generation of hepatocytes and development of the colon^71, 72^. Finally, to identify the combination of TFs driving human preimplantation development, we clustered regulon activity within lineages and ordered cells by pseudotime (**Fig. 2f**), revealing lineage-specific, regulatory states represented by 10 modules that comprise temporally distinct on/off-switches of TFs. In sum, these data predict regulators of cell identity and combinations of TFs that form the basis of the regulatory programs, cell state transitions and cellular identity in human preimplantation embryogenesis. We next investigated in depth whether aneuploidy induces changes in gene expression and gene regulatory programs, and whether these can be linked to cellular dysfunction in development.

### Gene dosage effects in expression vary with the nature of aneuploidy and the developmental stage

Although the transcriptomes of most genomically abnormal cells aligned with the euploid cells following clustering (**Extended Data Fig. 2b**), this does not rule out transcriptional and phenotypic changes resulting from (segmental) aneuploidy. To scrutinise any perturbations in gene expression, we first studied how gene expression varies with gene DNA copy number –i.e. ‘direct gene dosage effects’ (DGDE)– across preimplantation embryo development. Groups of single cells in the same developmental timepoint were defined by quantized transcriptome-pseudotime bins (**Fig. 3a**), rather than by embryonic day of development, as cells within the same embryo are not necessarily synchronised. Hence, these pseudotime bins offer an unbiased molecular-based classification of cellular developmental states or timepoints. Then, by comparing the expression levels of genes within DNA losses and gains relative to their euploid states in pseudotime-matched cells (**Fig. 3a**), and fitting a hierarchical Bayesian model on the observed deviations, we were able to decompose the observed DGDEs (**Fig. 3b**) into a pseudotime-invariable term (i.e. the average DGDE provided by a DNA gain or loss across the dataset) and a pseudotime-dependent term (i.e. the change in DGDE size across the different preimplantation developmental stages) (**Fig. 3b**). Importantly, the G&T-seq data, providing DNA-and RNA-readouts of the same single cells, allowed us to model DGDEs unambiguously, avoiding circular reasoning like inferring the copy number of a set of genes from scRNA-seq data and then modelling their direct gene dosage effects from the same scRNA-seq data^73^. For the pseudotime-invariable term, posterior distributions show positive effect sizes for DNA gains, and negative effect sizes for DNA losses (**Fig. 3b**), reflecting on average increased and decreased gene expression levels for genes hit by DNA gains and losses, respectively, as expected. However, for the pseudotime-dependent term, larger effect sizes around the pre-morula stage (i.e. Q1 pseudotime bin) were observed followed by a decrease in effect sizes around the early blastocyst stage (i.e. Q4 bin), and again increased effect sizes at ICM differentiation (**Fig. 3a-b**), suggesting that the magnitude of the DGDE changes with embryonic stage and is largely coinciding with intronic read ratios, which provide a proxy for the level of active transcription (**Fig. 3a**). Before EGA, minor DGDEs were detected, which, together with an intronic read ratio of >0 at those stages, might be indicative of a minor transcriptional wave before E3^74^. Furthermore, the magnitude of the increase in gene expression as a result of DNA gains was generally higher than that of the decrease in gene expression in DNA losses (**Fig. 3a-b**), suggesting that changes in DNA copy number do not result in strictly proportional transcript-level effects.

The observed DGDEs then led us to optimise and benchmark the inference of (segmental) aneuploidies in embryos from scRNA-seq data, using the DNA-sequences of the same single cells as ground truth (**Methods, Extended Data Fig. 5, Extended Data Table 3-4**). This in turn allowed us to include previously published datasets^45^, yielding 71% of cells (n = 2105) being classified as aneuploid and 88% of all embryos (n = 196) containing at least one aneuploid cell (**Fig. 4a-b**, **Extended Data Fig. 6**, **Extended Data Table 5**), and to determine gene-specific DGDEs. After EGA, we detected significant up-or down-regulation of expression for 556 and 428 genes out of 2325 and 2168 in DNA copy number gains and losses, respectively (Wilcoxon rank-sum test FDR corrected p-value < 0.001). Amongst the 100 most dosage sensitive genes (**Fig. 3c**), we identified genes involved in key cellular functions, including transcriptional regulation, RNA processing, translation, ribosomal components and regulators, protein degradation, autophagy and apoptosis, cell cycle regulation, DNA replication and proliferation and mitochondrion-related genes (**Extended Data Table 6**). Finally, the overall DGDE-fold changes for genes with a significant DGDE (adj p<0.05) in DNA losses and gains were 0.8 and 1.2, respectively (**Fig. 3d**).

**Figure 5.**
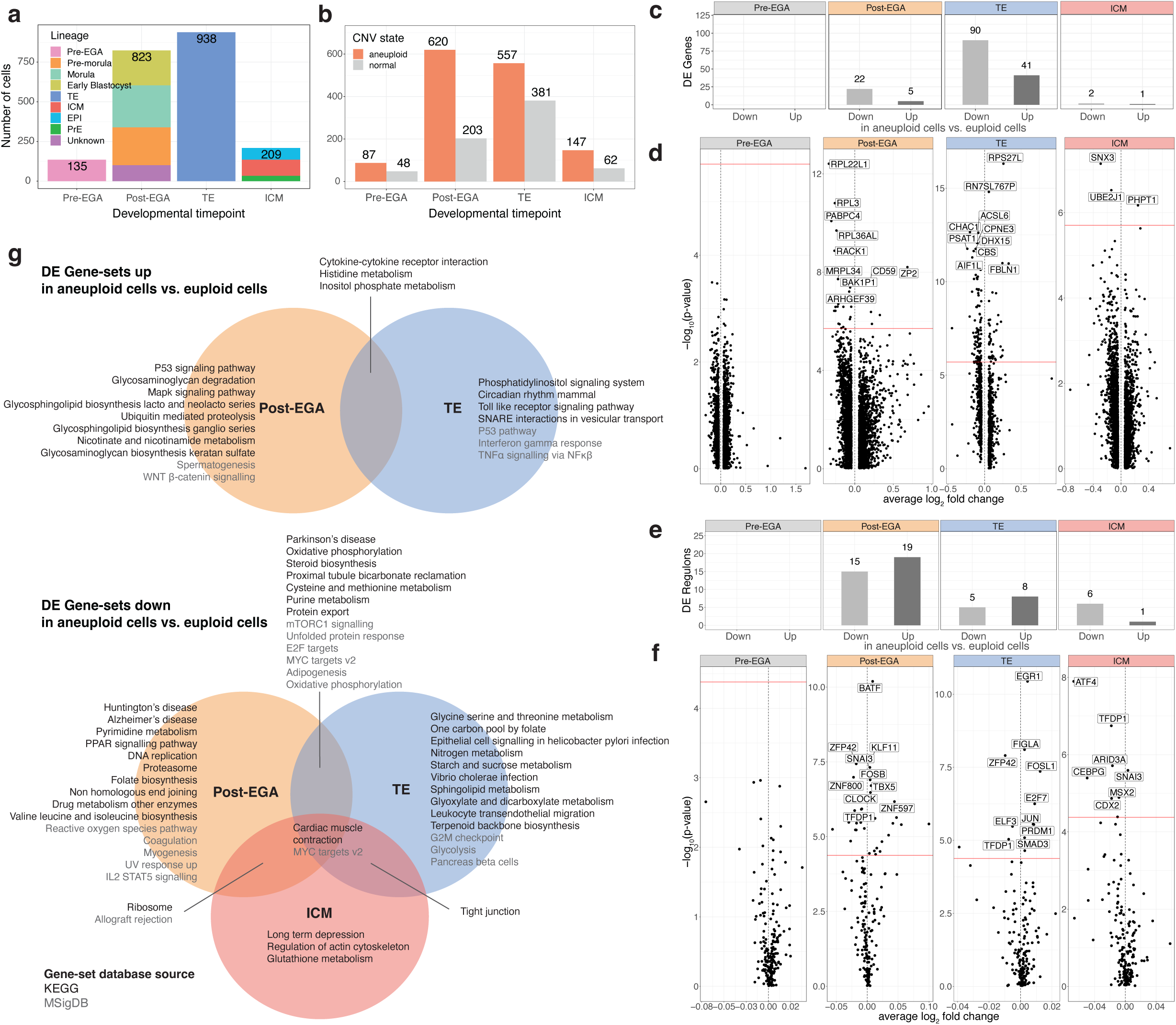
Differential expression of genes (DGE), regulons (DRE) and gene sets (DGSE) between euploid and aneuploid cells in different embryonic lineages. **a.** Lineage groups used for euploid-aneuploid cells differential expression analyses **b.** Number of normal and aneuploid cells per lineage group. **c,e .** Number of upregulated and downregulated genes and regulons in aneuploid cells in respect to euploid cells of each lineage group. **d,f.** Volcano plots showing downregulated (left) and upregulated (right) genes and regulons in aneuploid cells for each lineage group. The red line indicates the threshold p-value. The top 10 significant genes and regulons are labelled. **g.** Upregulated and downregulated KEGG and MSigD gene sets in aneuploid cells for each lineage group.

In summary, although (segmental) aneuploidies result in direct gene dosage effects, their magnitude depends on the nature of the aneuploidy (higher for gains than losses) and the embryo’s developmental stage (stronger at E3 and E5-6 than other stages). These DGDEs enable the inference of CNVs, with best performance post-EGA, and vary by gene.

### Copy-number aberrant genes coding for TFs perturb TF-target gene expression

TF genes, when deleted or duplicated, may in turn affect the expression of their euploid target genes, a process which we termed the ‘indirect gene dosage effect’ (IGDE) of a TF-gene, and perturb the gene regulatory network. We developed a method for single-cell IGDE analysis (**Methods**), and detected significant IGDEs for 50 TFs following EGA (Fisher’s combined FDR corrected p-value < 0.01, n = 1257) in at least one pseudotime bin (**Fig. 3e-f**). Importantly, this includes perturbations of regulons known to be key for embryo development, like GATA3^53^. Additionally, inverse IGDEs were also detected, where shifts in the expression of target genes showed lower or higher expression for gains or losses of the upstream TF gene, respectively. For example, *CTCF* showed an inverse IGDE suggesting it can act as a transcriptional repressor^75–79^. Overall, this demonstrates how the transcriptional effects of CIN in the preimplantation embryo can be spread across the transcriptome by direct and GRN-mediated dosage effects.

### Effect of aneuploidies on developmental progression in preimplantation embryos

Given that aneuploidies may result in direct and indirect gene dosage effects, we next investigated their impact on lineage specification and developmental progression. First, we investigated the physical age of the embryonic cells –i.e., the day of embryo development since fertilisation– relative to their molecular pseudotime age on the transcriptional trajectory. Importantly, this revealed that euploid cells, particularly at E3, E4, E5 and E7, tended to have progressed further in development, while aneuploid cells typically lagged in pseudotime (**Fig. 4c-d, Extended Data Table 7**), although few appeared to have accelerated. Next, we determined whether, within the same embryo, the cells that are delayed in developmental progression have a more abnormal genome than their sister cells. For each embryo, we computed how transcriptome-pseudotime ages of its constituent single cells correlated with the fractions of the genome that were aberrant in those same single cells (**Fig. 4e-f, Extended Data Fig. 7**). Eleven of 141 E3-E7 embryos (8%) showed a significant negative correlation, supporting a developmental delay of genomically aberrant cells within an embryo (**Fig. 4e-f**). Although a majority of other embryos showed the same tendency (**Extended Data Fig. 7**), three embryos showed a significant positive correlation, two of which carried putative meiotic abnormalities (**Extended Data Fig. 6**).

In conclusion, we show that acquired (segmental) aneuploidies lead to intra-embryonic cellular heterogeneity in molecularly-defined developmental age, whereby cellular differences in genomic aberrations generally correlate with developmental delay, and in rare instances a developmental acceleration.

### Cell fitness and competition in human preimplantation development

In order to reconcile the direct and indirect gene dosage effects with the developmental delay observed in aneuploid cells, we next investigated cellular responses to (segmental) aneuploidy by differential gene, regulon, and gene-set expression analyses (abbreviated as DGE, DRE and DGSE, respectively) between euploid and aneuploid cells in pre-EGA, post-EGA, ICM and TE (**Fig. 5a-b**). Except for pre-EGA cells, all lineages showed significant differences (**Fig. 5c-g**).

In line with their developmental delay phenotype, gene expression signatures indicated aneuploid cells to be less fit across all cell lineages following embryonic genome activation (**Fig. 5c-g**). In post-EGA aneuploid cells, genes involved in nucleotide metabolism (e.g. *UCK2*^80^), mRNA stability and processing (e.g. *LRPPRC*^81^), ribosome biogenesis, ribosomal function or translation (e.g. *RPS19BP1*^82^), respiratory chain assembly (e.g. *NUBPL*^83^) and mitochondrial ribosomal function (e.g. *MRPS12*^84^) (**Fig. 5d**, **Extended data Table 8**) were down-regulated. Using DGSE, we found the ribosome gene set to be most significantly down-regulated in aneuploid cells in addition to other cell function, growth and metabolism-related gene sets, including the mTOR signalling pathway^85, 86^ and MYC target genes^87^. In contrast, the p53 signalling pathway, amongst others, was up-regulated (**Fig. 5g**). Furthermore, DRE identified anti-proliferative or pro-apoptotic regulons with increased activity in aneuploid cells, such as the p53 activator *EGR1*^88^, the tumour suppressor *KLF10*^89^ and *BATF,* an inhibitor of AP-1 –the master regulator of cell growth and proliferation^90, 91^– whereas regulons with decreased activity are involved in proliferation (e.g. *TFDP1*^92^), metabolism (e.g. *PPARG*^93^) or have essential functions in pluripotency and embryo development (e.g. *FOXK2*^94^, *POU5F1*^61^, *CDX2*, *ZFP42*^95^, *ARID3A*^96^) (**Fig. 5f**). Additionally, the expression of genes and the activity of regulons with oocyte-related functions are increased in aneuploid post-EGA cells (**Fig. 5d,f, Extended Data Table 8**). Similar to post-EGA, the predominantly altered gene-expression signatures in ICM and TE aneuploid cells are indicative of a developmental delay phenotype (**Fig. 5d,f,g, Extended Data Table 8**). Both the mTOR-and Hippo-YAP/TAZ signalling pathways are down-regulated in post-EGA and TE aneuploid cells (**Fig. 5d,f,g, Extended Data Table 8**), and have previously been implicated in reduced cell fitness and competition events in the mouse embryo^86, 97–100^. In sum, we detected gene-expression signatures in aneuploid cells across all lineages after EGA that are consistent with an aneuploidy-induced developmental delay and reduced cell fitness.

The unfolded protein response (UPR), which is triggered by increased unfolded or misfolded proteins in the endoplasmic reticulum (ER), represents a survival response in cells to restore ER homeostasis^101^ and is often activated in aneuploid cells of other models, together with the ubiquitin-proteasome (UPP) or autophagy-lysosomal (ALP) protein degradation machineries^102^. However, except for the upregulation of the *DDIT3* regulon post-EGA, which is a pro-apoptotic transcription factor^103^, we found that the UPR salvage pathway that promotes cell survival is suppressed in aneuploid cells of the human preimplantation embryo. Indeed, the UPR gene set is downregulated in aneuploid cells of post-EGA and TE lineages, including its key components^101^ such as the *ATF4* regulon in post-EGA and ICM, the *ATF6* regulon in post-EGA, the MYC target gene-set in all lineages, and the *CEBPG* regulon^104^ in ICM (**Fig. 5d,f,g**). Furthermore, *UBE2J1*, which catalyses the covalent attachment of ubiquitin to proteins and which functions in the selective degradation of misfolded membrane proteins from the endoplasmic reticulum as part of the UPP degradation machinery^105^, is found significantly down-regulated in the aneuploid ICM cell population (**Fig. 5d, Extended Data Table 8**). Also, ALP-related features^106^ were found decreased in aneuploid post-EGA and TE cells, including gene sets involved in neurodegenerative disorders with proteotoxic stress (e.g. Alzheimer’s, Parkinson’s and Huntington’s), the mTOR pathway, the AP-1 transcription factor complex in post-EGA and the sphingolipid gene-set in TE (**Fig 5f-g**). Hence, our data also indicates that important components of cellular proteostasis are suppressed in aneuploid cells of all lineages, likely further contributing to their cell unfitness.

Finally, in aneuploid TE cells gene expression signatures suggestive for both pro-and anti-proliferative cellular phenotypes were detected (**Extended Data Table 8**). *CDKN1A*, for instance, which promotes cell cycle arrest^107^, as well as its inhibitory target CCND1, which promotes cell cycle progression^108^, were found to be significantly increased in aneuploid TE cells (**Fig. 5d**, **Extended Data Table 8**). Also, a significant number of genes expressed in the placenta or related to implantation were found perturbed in comparison to euploid TE cells (**Fig. 5d,f,g**, **Extended Data Table 8**), suggesting that aneuploid TE cells may affect implantation features^31, 109^.

Importantly, our gene expression analyses indicate that aneuploid cells in human preimplantation embryos develop an unfit cellular phenotype post-EGA onwards, corroborating their developmental delay in transcriptomic pseudotime.

## DISCUSSION

Studies investigating gene expression in human preimplantation embryos at single-cell level are often based on transcriptome analysis only and frequently overlook their extraordinary susceptibility to chromosome instability^42, 45, 51, 110, 111^. However, the genomically unstable nature of human preimplantation embryos, as well as the developmental transcriptional dynamics, mandate a single-cell multi-omics approach to develop an understanding of the (an)euploid cell biology in human preimplantation embryos. Using single-cell genome-and-transcriptome sequencing on all single cells from the same embryo, we show that (segmental) aneuploidy significantly perturbs gene expression and its regulation, and in turn impacts cell fitness, leading to cell-competition signatures within the developing human embryo.

Integrated DNA-and RNA-seq analyses demonstrate that direct gene dosage effects initiate in human embryonic cells at EGA, with effect sizes depending on developmental stages thereafter. Surprisingly, these DGDEs are generally milder than expected from the DNA copy number of the affected genes. In one model, the size of the DGDE for genes within a newly acquired DNA-loss will be underestimated because of the transcripts of the same genes inherited from the euploid precursor cell. However, in an alternative model, dosage compensation may also lead to milder DGDEs for DNA losses^112^ through increased expression of genes on the remaining chromosome, similar to the increase of transcription of genes on the single X chromosome in male *D. melanogaster*^113, 114^ or in mouse embryonic stem cells with monosomy X^115^. Our findings are possibly concordant with such a mechanism since the fold-change in expression of genes in regions with DNA-losses is only ∼0.8 instead of the expected 0.5. This is in fact similar to the ∼0.7-0.6 fold-change reported in a recent study on dosage compensation in monosomic non-transformed human cell lines^116^. Furthermore, in various somatic cell types, gene dosage compensation has been documented for viable human trisomies^117, 118^, whereby a significant fraction of genes on the 3-copy chromosome is down-regulated towards their euploid expression level^119^. In accordance with this, we find that the fold-change in expression for duplicated genes in general remains well below the expected 1.5 despite the acquired extra allele, indicating also that dosage compensation mechanisms may already be operative during early human development.

In gene expression regulation, TFs are often encoded on a different chromosome from their target genes^120^, raising the question whether GRNs become dysregulated if their cognate TF is hit by a (segmental) aneuploidy. We demonstrate that the expression of euploid target genes within a regulon can be significantly perturbed by an upstream copy-number aberrant TF within single cells of the mosaic aneuploid-euploid embryo. Depending on when and which TFs are hit by (segmental) aneuploidy during preimplantation development, these can include GRN-expression perturbations of TFs important for early development, like GATA3 in this study, which could impact downstream lineage commitment of those cells. Indeed, previous studies that artificially perturbed the expression of TFs in human embryos^53, 61, 63^, including the down-regulation of GATA3 in the human morula stage^53^, showed how this can lead to developmental failure of the embryo. The CIN-driven GRN perturbations we identified in the human preimplantation embryo and their effects on embryogenesis will not only depend on the timing of the acquired copy-number aberrant TF (whereby pre-EGA events may affect the developmental potential of a larger fraction of cells in the embryo), but also on the loss-or gain-of-function nature of the genetic aberration and on the embryonic cell type it is acquired in. As a guideline to further study these indirect gene dosage effects in human embryogenesis, this study also provides a comprehensive resource of all TFs and GRNs that are activated at specific stages over the course of human preimplantation development, and uncovers both known and as yet uncharacterised TFs with likely important roles in early embryogenesis.

Next, we investigated how aneuploidy-induced direct and indirect gene-expression dosage effects impact developmental progression of the aneuploid cells in the embryo. First, we compared the cells’ post-fertilization ages to their pseudotime ages on the transcriptomic trajectory. Remarkably, aneuploid cells often lagged in their development in comparison to euploid cells, suggesting a decline in cell fitness with the acquisition of genomic anomalies. This is indeed endorsed by gene-expression signatures indicating impairment of (mito)ribosome biogenesis, translation, energy homeostasis and proliferation within aneuploid cells following EGA. This gene-expression signature is further corroborated by other studies that investigated the consequences of aneuploidies in primarily blastocyst-stage human embryos using either single-cell unimodal RNA-seq^35^ or multi-cell gene expression studies^32, 36^, and in other eukaryotic model systems^25, 27^. Additionally, aneuploid cells can be depleted from all embryonic germ layers in later stages of development^31, 121^. Together, these observations support a model whereby the aneuploid cells with declined fitness become the losers and the euploid cells the winners in a cell-competition game ensuing post EGA and which is instigated by chromosome instability. Cell competition is known as a fitness-sensing and quality-control mechanism that can eliminate cells which, although viable, are less fit than their neighbours^122^, suggesting a possible way by which mosaic embryos may lead to healthy offspring^13, 19, 20, 30^.

We then investigated what molecular mechanism could trigger the onset of the cell competition. Recently, ribosomal protein (Rp) genes have been shown to act as sensors of (segmental) aneuploidy in *Drosophila*, whereby Rp gene dosage sparks cell competition via the Rps11-Xrp1-pathway, leading to the elimination of (segmental) aneuploid cells by surrounding diploid cells^123–125^. Controlled removal of large chromosomal segments indicated that cells which retained their *Rp* genes survived, while those that lost *Rp* genes usually died though intriguingly only when surrounded by normal cells. Similar to *Drosophila*, the 80 human ribosomal proteins are mostly encoded by single copy genes that lie scattered throughout the human genome^126, 127^, suggesting these could similarly act as sensors of aneuploidy in human cells. In line with these observations, the large (segmental) aneuploidies in the human embryo affect *RP* gene expression, with *RP* genes being amongst the most significantly affected by direct dosage effects in our study, which in turn, may perturb ribosome biogenesis^128^ and instigate cell competition downstream. The transcription factor Xrp1 was recently shown to be a master regulator of cell competition instigation in *Drosophila*^129^. Elevated expression of Xrp1 in Rp mutant *Drosophila* cells restricts translation, cellular growth, and mosaic organismal development^123^. Although there is no direct homolog of Xrp1 in humans, the human bZip domain protein most similar to the Drosophila Xrp1 is DNA Damage Induced Transcript 3 (DDIT3 or CHOP)^130^, and also mediates multiple functions of p53^131^, which we found upregulated at the regulon level in aneuploid cells post-EGA, aligning with the cell competition model of *Drosophila*. DDIT3 is known as a multifunctional transcription factor in the ER unfolded protein response, and fulfils an essential role in inducing cell cycle arrest and apoptosis when the UPR salvage pathway fails. Although the UPR is known to be activated in response to aneuploidy^132^, we observed a significant down-regulation of key regulons involved in UPR in aneuploid cells in the embryo, including down-regulation of ATF4 in post-EGA and ICM, and ATF6 in post-EGA, as well as a down-regulation of the UPR hallmark gene-set and other proteotoxic stress related gene-sets (i.e. Huntington’s, Alzheimer’s or Parkinson’s diseases) in post-EGA and TE. This scenario is concordant with the Rp mutant model of cell competition in *Drosophila*. In this model, the UPR genes are not upregulated despite the presence of the ER stress signatures^129^. In *Drosophila* Rp mutants, up-regulation of UPR genes is observed only if Xrp1 is repressed, leading to the hypothesis that Xrp1 instigates cell competition by dampening UPR and thus, avoiding that the ’loser cells’ get repaired, and instead get substituted by adjacent wild type proliferating cells^129^. This is in line with our findings in the mosaic human embryo. We propose that downregulation of UPR, reduced translation and upregulation of DDIT3 provide a mechanism for the elimination of aneuploid cells from the mosaic embryo.

Other initiators of cell competition have been identified. For instance, mitochondrial dysfunction triggers cell competition in mouse epiblast cells^133^ and appears to be common to a range of different loser cell types^134–136^. We show that aneuploid cells in the human embryo also acquire dysregulation of mitochondrial gene expression after EGA in all lineages, and share other features of loser cells as described for normal mouse epiblast development, such as down-regulation of mTOR pathway^86^, MYC targets and pluripotency genes, as well as YAP inactivation and up-regulation of the p53 ^pathway^^97–100^. In summary, our results suggest that cell competition mechanisms eliminating unfit _epiblast cells_ in mouse embryos –even though they are unrelated to aneuploidy– are likely also triggered by (segmental) aneuploidy during human preimplantation development.

In summary, our multi-modal analyses reveal novel insights into the first week of human development. We provide a unique comprehensive resource of the TFs and gene regulatory networks driving preimplantation embryo development, and how this is perturbed with CIN, and how this, in turn, may lead to cell competition within the mosaic human embryo.

## MATERIAL AND METHODS

### Embryo source

Cryopreserved embryos donated for research were obtained from couples that underwent IVF with or without PGT at Leuven University Fertility Center (LUFC). Embryos used for this study were cryopreserved either at zygote stage (E1 -with 2 pronuclei and 2 polar bodies), at cleavage stage (E3 - if sufficient morphological quality^137^) or at blastocyst stage (E5 or E6 - if sufficient morphological quality^138^) and diagnosed as affected and/or aneuploid for a viable trisomy by comprehensive PGT^139, 140^. Embryos cryopreserved at the zygote or cleavage stage were supernumerary after IVF treatment. For those, couples signed an informed consent to allow their use for research after the legal storage period of 5 years. Embryos cryopreserved at the blastocyst stage were non-transferable after comprehensive PGT. For those, couples signed an informed consent to allow their use for this specific research during treatment. A total of 123 zygotes, 88 cleavage stage embryos and 14 blastocysts were thawed, of which 68(55%), 43(49%) and 13(93%) survived, respectively.

### Embryo thawing and culture

Cryopreserved embryos were warmed following Vit Kit®-Thaw (Irvine Scientific) manufacturer’s instructions. After thawing, each embryo was cultured between 5h to 48h in a well of a Universal GPS® dish (LifeGlobal) containing 40µl of GM501 medium (Gynemed) under mineral oil (Gynemed) at 37°C, 6% CO2, pH 7.25–7.35 in a time lapse incubator (ASTEC Penguin incubator). Zygotes were cultured to obtain E1, E2 and E3 embryos; cleavage stage, to obtain E3, E4 and E5 embryos; and blastocysts, to obtain E5, E6 and E7 embryos. Due to the requirements of manual single-cell dissociation, warming and culture of embryos was planned so that a maximum of 8 E1-2, 4 E3-4, 1 E5-7 were dissociated on the same day.

### Embryo dissociation and collection of single cells

For dissociation, each embryo was removed from the culture media and placed in 40µl of prewarmed Ca^2+^ Mg^2+^ free media (PGD biopsy Medium, LifeGlobal). Subsequently, zona pellucida was removed by placing the embryo in 20µl of pre-warmed Acid Tyrode at 37°C (pH 2.4 Sigma-Aldrich). After washing through clean prewarmed Ca^2+^ Mg^2+^ free media, the embryo was dissociated into single cells using Stripper® pipette and capillaries (CooperSurgical) of several decreasing diameters depending on the embryo stage: 175µm, 135µm, 75µm and 50µm. Prior to dissociation, only embryos at blastocyst stage were placed in 100µl 0.05% Trypsin EDTA (Thermo Fisher Scientific) for 10 min at 37°C. Each isolated single cell was washed in 3 fresh 7µl drops of PBS 1%PVP (Sigma-Aldrich) and placed in a well of 96-well plate containing 2,5µl of RLT plus buffer (Qiagen). At the end of cell collection, the plate was centrifuged for 1 min at 1000g and stored at −80°C until further processing. Cells of the HapMap cell line GM12878 were used as single-cell and multi-cell controls which were added to each collection plate together with negative controls.

### Separation, amplification, library preparation and sequencing of DNA and RNA from single cells

Single cells were processed according to G&T-seq protocol ^141^. For the mechanical separation of DNA and RNA, we used a liquid handling robot (Microlab STAR Plus, Hamilton). Each 96-well plate, containing isolated cells in 2,5µl RLT plus buffer supplemented with 1µl of ERCCs (Life Technologies), was placed in a predefined location of the robot deck together with (1) a plate containing 10µl per well of Dynabeads® MyOne^TM^ Streptavidin C1 (Thermo Fisher Scientific) bound to biotinylated poly-dT oligos containing the SmartSeq2 primer sequence ‘5BioTinTEG/ - AAGCAGTGGTATCAACGCAGAGTACTTTTTTTTTTTTTTTTTTTTTTTTTTTTTTVN’ (IDT) – to capture poly-A mRNAs, (2) a plate containing 25µl per well of G&T-wash buffer – to wash away the DNA from the cell lysate supernatant, and (3) an empty DNA destination plate. First, the cell lysate was mixed and incubated for 20 min with biotinylated poly-dT beads. Then, beads bound to mRNA were pulled down using a low elution magnet (Alpaqua) for 2 min and, subsequently, supernatant was transferred to the DNA destination plate. Lastly, beads were washed 2 times with 10µl of G&T wash buffer, i.e. mixed, incubated 5 min and pulled down for 2 min, and supernatant was transferred to the DNA destination plate. To avoid loss of material, the same set of Axygen® FXF-384-XL-R-S tips (VWR) were used for the entire separation protocol and were rinsed with 5ul of G&T wash buffer at the end of the separation step. The DNA destination plate containing 37,5µl of G&T wash buffer was centrifuged 1min 1000g and stored −80°C. The plate containing only beads bound to poly-A mRNA was immediately processed following adapted SmartSeq2 protocol ^142^ with 20 PCR cycles. Amplified single-cell cDNA was purified with 1:1 ratio of Agencourt AMPure XP beads (Analis) or CleanNGS beads (GC Biotech), washed with 80% ethanol and eluted in water. Concentration and fragment size of amplified cDNA was measured with QuantiFluor® dsDNA (Promega) and Fragment Analyzer (Advanced Analytical) High Sensitivity kit DNF-486 or 487, respectively. Prior to whole genome amplification (WGA), the single-cell DNA in 37,5µl of G&T wash buffer was purified using the liquid handling robot (Microlab STAR Plus, Hamilton) where 25,5µl of Agencourt AMPure XP beads (Analis) or CleanNGS beads (GC Biotech) were added to each well. The mixture was incubated for 20 min, the beads were pulled down on a low elution magnet for 20 min and were washed twice with 50 µl 80% ethanol. The plate containing dried beads bound to single-cell DNA was extracted from the robot and processed following PicoPLEX® (Sopachem/Takara) protocol with half volumes as described in Macaulay *et. al.* (2016) ^141^. Amplified single-cell gDNA was purified and quantified as described above for cDNA. Libraries from amplified cDNA and gDNA were prepared following the Nextera XT (Illumina) protocol with quarter volumes. Libraries were pooled and sequenced maximum 96 samples single-end 50 on one lane of a HiSeq4000 or HiSeq2500 Illumina sequencer aiming at 1 million reads per sample. DNA libraries were aligned using BWA (version 0.7.17 https://arxiv.org/abs/1303.3997) in “mem” mode, using default parametrization, to the GRCh37 human reference genome build. PCR duplicates were removed using Picard (version 2.19).

### Single-cell copy number variant detection

Estimation of DNA copy-number variation by focal read depth analysis was performed as previously described^141^ with minor modifications. In brief, single-end sequencing reads were demultiplexed and aligned to the GRCh37 human reference genome with BWA^143^ (Version 0.7.17-r1188 https://arxiv.org/abs/1303.3997) in “mem” mode, using default parametrization). PCR-duplicates were removed using Picard (Version 2.19.0) (http://broadinstitute.github.io/picard/). Genomic bins were determined as non-overlapping 250,000 uniquely mappable positions. For each bin, LogR was calculated as the read count of that given bin divided by the average read count of the bins genome-wide plus one. LogR values were normalised according to the median LogR genome-wide and correction for %GC content was performed using a Loess fit (with a span of 0.3) in R. The corrected LogR values were segmented into regions with a similar value using piecewise constant fitting (PCF) with the penalty parameter, g, set to 15. Integer DNA copy numbers (CN) were estimated as 2^logR*y, with the average ploidy of the cell, y=2. Chromosome 19 which presented a higher loss rate due to its known high %GC content^144^ (**Extended Data Fig. 1a**), was not considered for downstream DNA-based CNV analysis such as cell classification and inferCNV benchmarking. The initial quality filter for single-cell DNA libraries was set at MAPD score < 1.5. However, after visual inspection of all genome-wide CNV profiles, 3 cells (i.e., e2_2, e11_8, e11_5) were retained due to their concordant result with the other cells from the same embryo and 2 cells were eliminated due to the incorrect logR segmentation due (i.e., e33_3, e49_20), resulting in 295 out of 434 cells passing these quality control criteria. Per cell, a genomic read coverage of 0.15x was obtained, sufficient for DNA copy number calling (**Extended data Fig. 1a-b**).

### Single-cell RNA-seq data pre-processing

RNA-seq libraries of the G&T-seq processed cells received a median of 1.47M uniquely mapped reads, detecting an average of 10354 genes per cell (**Extended Data Fig. 2a**). Cells with fewer than 3000 expressed genes, a mitochondrial read fraction higher than 30% or an ERCC fraction higher than 20% were discarded from further analysis, resulting in 576 cells out of 827 passing these quality control thresholds. Raw scRNA-seq data from ^45^ were obtained from EMBL-EBI Array Express (E-MTAB-3929). All reads from the G&T-seq RNA libraries and the Petropoulos dataset were mapped to the reference genome (GRCh37.75) using STAR aligner (v2.5.2b) with GENCODE release 19 gene annotations. Cell-wise counts were obtained using HTSeq (v0.12.4). These gene-wise counts from the two experiments were then merged into a single count matrix. Counts were then normalised and batch corrected using the ‘Integration and Label Transfer’ standard workflow within Seurat (3.0.2). InferCNV v1.3.3 was used to derive copy number estimates from the scRNA-seq data^73^. Briefly, cells were split into ten pseudotime bins with an equal number of cells in each and processed using default parameters apart from setting cutoff = 1, ref_subtract_use_mean_bounds = F, analysis_mode = “cell”, and HMM_report_by = “cell”. Chromosomes X, Y and Mt were excluded and the average signal across all cells in a pseudotime bin were used to define the baseline. The 6-state Hidden-Markov Model was used to assign copy number states.

### Regulon analysis

SCENIC^46^ was performed on the integrated data using pySCENIC (0.9.15). pySCENIC was run 5 separate times on the data. From the 5 runs, ‘unstable’ regulons which only appear in one run were removed. The highest AUC value among regulons duplicated in more than one run was kept to build the final AUC matrix. Following SCENIC, regulons were ranked for their specificity within each lineage using the method developed by Suo et al. (2018). Pseudotime was performed using slingshot (1.0.0). To construct individual networks (**Extended Data Fig. 4**) for each lineage we first ran the binarization step of SCENIC (pySCENIC 0.9.15) on the SCENIC generated data in order to establish an on/off status for each regulon in each cell. We then pruned connections (between TFs and target genes) to only those which appear in all 5 runs of SCENIC. Each network was constructed from cells belonging to the specified lineage and with regulons which appear in at least 80% of all cells of said lineage.

### Trajectory analysis

To determine temporally expressed genes we applied a general additive model (gam 1.16.1) to the normalised count data. Here cells were split into two sets determined by the two slingshot pseudotime trajectories (visualised as two lines in **Fig. 2d**) leading to either a TE or ICM/EPI/PrE end point.

### Cell typing

To assign cell type, cells were initially clustered based on the regulon AUC matrix, clustree (0.4.0). Here we identified which clusters likely belong to each lineage, based on expression of known marker genes (NANOG for Epiblast, SOX17 for primitive endoderm, GATA3 for trophectoderm, etc.). We then performed differential gene expression analysis between the selected clusters (Seurat 3.0.2), and selected genes which were highly up-regulated and genes which were not present within each cluster to build an annotation file. This annotation file was supplied to the automatic cell type classification program, Garnett (0.1.4) along with the raw counts matrix to perform cell type classification (**Fig. 2c, Extended Data Fig. 3e**). All TE subtypes were pooled following classification.

### Direct gene dosage effect analysis

To compare normal and abnormal copy number gene expression distributions, median normalised gene expression values were centred relative to the median gene expression of euploid cells in each pseudotime bin (**Fig. 2b**). For the detection of dosage effects, we restricted our analysis to genes that were expressed in at least 10 cells of the pseudotime bin and exhibited an euploid normalised median gene expression > 1.0. We then fit a Bayesian hierarchical linear model using STAN (v2.19.2) where *y* = *α* + *β_CN_C_i,j_* + *δ_CN,q_C_ij_q_j_* with y being the measured median-centred expression, *α* being the intercept representing the average median-centred expression of the euploid cells at E1, *β_CN_* representing the mean effect size per CN type as measured in *_Ci,j_* corresponding to the CN of gene *i* in cell *j. δ_CN,q_* represents the time-dependent effect size of each CN type for each pseudotime quantile q. Standard Gaussian priors with mean 0 and standard deviation 1 were chosen for *β_CN_* and *δ_CN,q_*, and posterior densities for these coefficients were estimated by running 4 Markov chains with 10000 iterations and 500 warmup sampling iterations with random parameter initiations. The total dosage effect was estimated as the sum of the time-invariant dosage effect size *β_CN_* and the time-dependent dosage effect size *δ_CN,q_* for each CN type.

We generated intronic and exonic read counts using the Velocyto python package (v0.17.17) and calculated the intronic read ratio as the ratio of intronic counts over total counts for each cell.

### Benchmarking of inferCNV with G&T-seq data in human embryos

We used the multi-omics G&T data to benchmark inferCNV-derived copy number calls. We directly evaluated this approach using the DNA-based copy number calls from the scG&T data as a ground truth. We defined base-wise true-positive and true-negative rates as the fraction of the DNA-seq based aneuploid or diploid segments, respectively, that are recapitulated by the scRNA-based InferCNV calls (**Extended Data Fig. 5a**). Calls made on the DNA-seq data were considered ground truth and segments were assigned as (i) true positive (TP) if both DNA and RNA copy number calls indicated a gain or if both indicated a loss; (ii) true negative (TN) if both indicated a copy number equal to 2 (note that both callers will miss copy neutral loss of heterozygosity events); (iii) false negative (FN) if the DNA data indicated a copy number aberration and the RNA did not; (iv) false positive (FP) if the RNA indicated a copy number aberration and the DNA did not (**Extended Data Fig. 5a**). Per-cell rates were calculated as the segment-length weighted rates. Aggregating (true) positive and negative calls across the cohort confirms that the accuracy of the RNA-based approach is similar across the genome (**Extended Data Fig. 5b**). As expected, due to maternally derived RNA, RNA-based inference fails to reveal DNA copy number pre-EGA, and the true-positive rate increased dramatically from a median of 0 for the first two pseudotime quantiles, which contain E1-E2 cells, to >0.5 for the subsequent ones, while the true negative rate dropped slightly from a median 0.97 to 0.87 (**Extended Data Fig. 5c**). Thus, the inferCNV approach can detect about 50% of aberrant copy number regions and more than 80% of normal genomic regions but only after EGA.

Chromosome arms (excluding the p-arms of the acrocentric chromosomes 13–15 and 21–22) were classified as aberrated when the fraction of that arm which had a copy number state ≠ 2 exceeded 60%. This threshold was optimised on the G&T data by computing true and false positive/negative rates as indicated above but at the arm level. Precision, recall and F1-scores were also calculated and indicated 60% of an arm as the optimal cutoff to designate it as aberrated. At this threshold, we observed optimal agreement between DNA and RNA-based arm-level copy number post-EGA and a precision and recall for cell classification of 0.71 and 0.77, respectively (**Extended Data Fig. 5d**). The same metrics were computed at the level of cell classification, confirming the 60% threshold (**Extended Data Fig. 5e**). A whole cell was considered aberrated when at least one arm was classified as such.

### Indirect gene dosage analysis

To calculate the dosage effect of aneuploid copy-number states of transcription factors on downstream euploid target genes, we first determined downstream target genes in for each TF’s regulon as described in the section “Regulon Analysis”. For each pseudotime quantile we tested for differences in the distributions of the median-centred expressions (as described under “Direct gene dosage effect analysis”) of target genes in cells with TF amplifications or deletions compared to euploid cells in that same pseudotime quantile. We only tested for TFs for which at least 10 cells were observed with either an amplification or a deletion of the TF in question within that quantile. To test for overall indirect gene dosage sensitivity we combined both amplification and deletion p-values using a Fisher’s Omnibus test and corrected for false discovery rate inflation due to multiple comparisons using the Benjamini-Hochberg method.

### Differential gene, regulon and gene-set expression analysis

Differentially expressed genes between euploid and aneuploid cells were determined by pseudotime quantile and by annotated lineage. These were computed using the FindMarkers function in Seurat using the “MAST” method with a logfc threshold of 0.05 and a min.pct threshold of 0.5. While testing for differentially expressed genes in the annotated lineages the pseudotime was given as a latent variable in the model. Multiple testing correction was performed by adjusting the p-values using the Benjamini-Hochberg method using the p.adjust function in R. Gene labels were plotted for genes with a −log10(pvalue) < 5.7, corresponding to the Bonferonni threshold for genome-wide significance with a significance threshold of 0.05 and the number of multiple comparisons set to 4 times the number of genes tested. For differential geneset analysis, genesets were obtained either through manual curation or from the KEGG Pathway annotations using the msigdbr R package (version 7.2.1). Regulon genesets were obtained using PySCENIC as described above. AUC values per geneset for each cell were computed using the AUCell R package (version 1.8.0). Differential geneset expression between aneuploid and euploid cells was determined per pseudotime quantile and annotated lineage by performing a Wilcoxon ranked-sum test on the AUC values. P-values were then adjusted for multiple comparisons using the Benjamini-Hochberg FDR-correction.

### Interpretation of differential gene, regulon and gene-set expression between euploid and aneuploid cells

To determine and classify relevant dysregulated functions we applied gene ontology analysis (GO) as well as manual literature curation for each gene in the context of embryonic development, aneuploidy, stress response, proliferation and apoptosis.

## Data availability

Data will be made available through EGA.

## Code availability

Code is available upon request.

## Ethics statement

This study was approved by the Ethics Committee Research University Hospital / KU Leuven (S58250, S65358) and by the Federal Commission for Embryo Research in Belgium (ADV_62, ADV_089).

## Acknowledgements

E.F.G. is a PhD fellow of the Research Foundation – Flanders (FWO). J.D. is a postdoctoral fellow of the FWO. Research in the lab of T.V. relating to this work is supported by the Research Foundation– Flanders (FWO; G081318N; G0C6120N; G088621N), the KU Leuven (SymBioSys - C14/18/092; C14/22/125; IDN/19/039), infrastructure grants (type 1 funding from the Hercules Foundation— AKUL/13/41; Foundation Against Cancer project 2015-143; and FWO I001818N) and European Union’s Horizon 2020 research and innovation program under grant agreement No 824110 – EASI-Genomics. Research in the Pasque lab is supported by The Research Foundation–Flanders (FWO) (Odysseus Return Grant G0F7716N to V.P., FWO grants G0C9320N and G0B4420N to V.P.), the KU Leuven Research Fund (BOFZAP starting grant StG/15/021BF to V.P., C1 grant C14/16/077 to V.P. and project financing).

## Author contributions

T.V. and V.P. conceived the study. E.F.G. performed embryo experiments with the help of A.K. A.S., J.C. and J.D. performed computational analysis with the help of J.C.H., R.V., M.V.d.H. and S.V., J.V.H., I.C.M. and T.V. developed and set-up automated G&T-seq, and basic analytical framework. S.D. and K.P. coordinated and supervised the warming of embryos donated for research. K.V. assisted in research management. E.F.G., A.S., J.C., J.D., V.P., J.C.H., R.V., J.R.V. and T.V. interpreted the data. E.F.G., A.S., J.C., J.D., V.P. and T.V. wrote the manuscript. All authors read and approved the manuscript.

