## Supplementary figures and images for "A multi-omics genome-and-transcriptome single-cell atlas of human preimplantation embryogenesis reveals the cellular and molecular impact of chromosome instability"

### Extended Data Figures1-4

#### Extended Data Figure 1

a

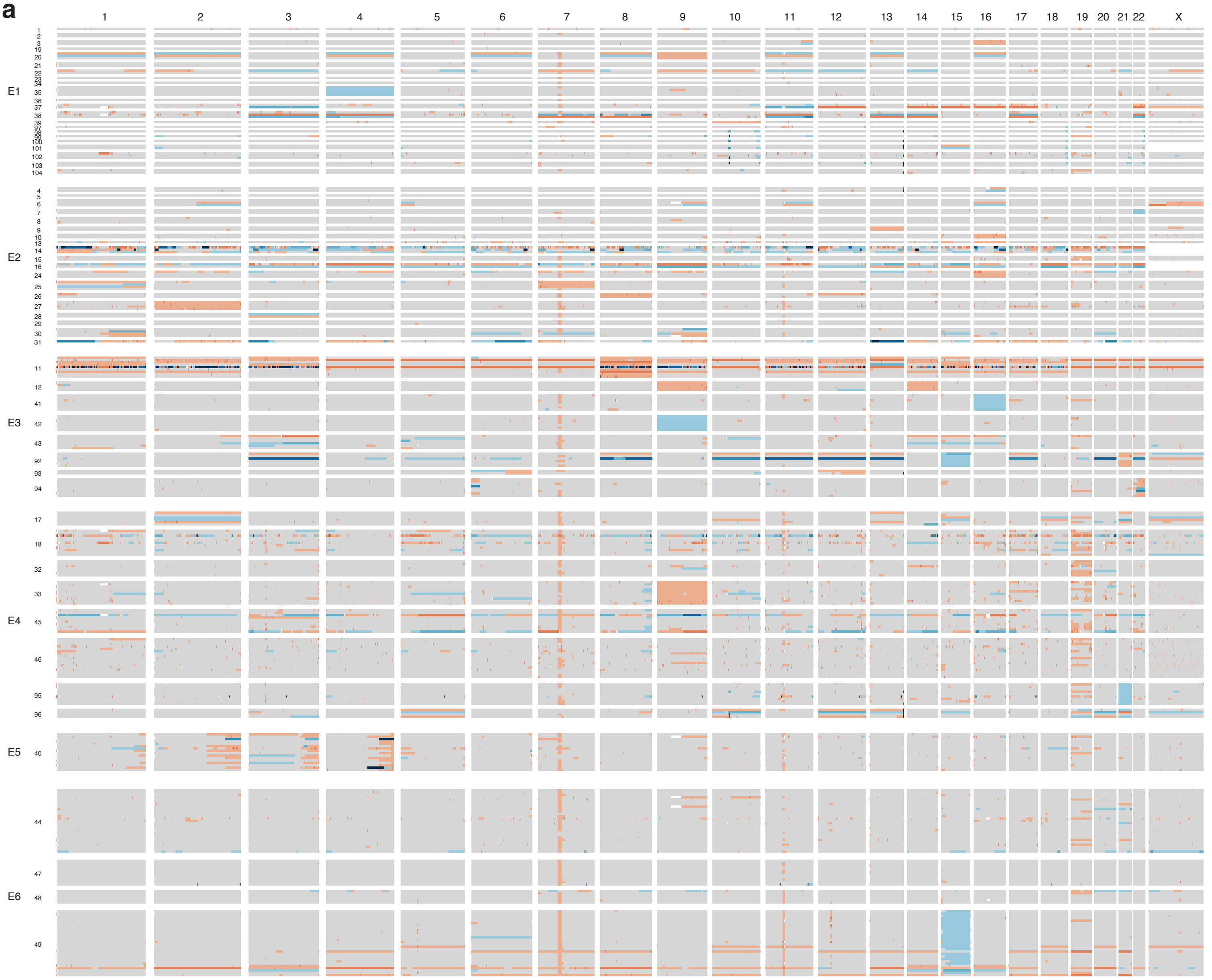


b

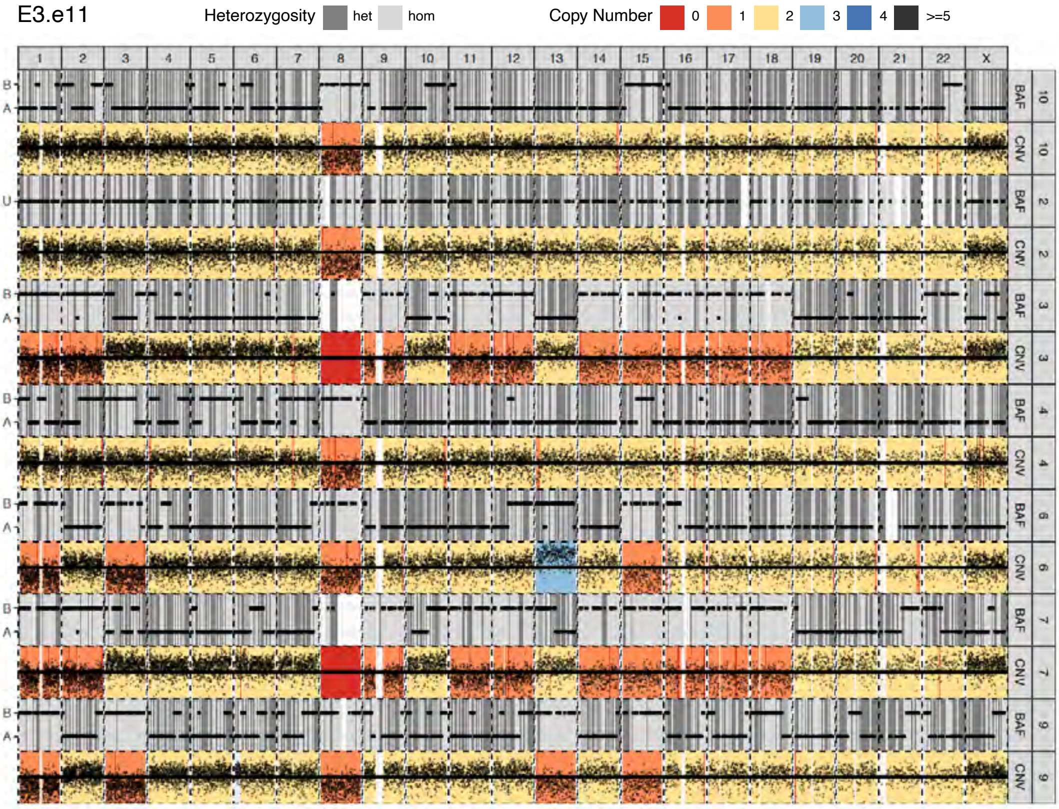


c

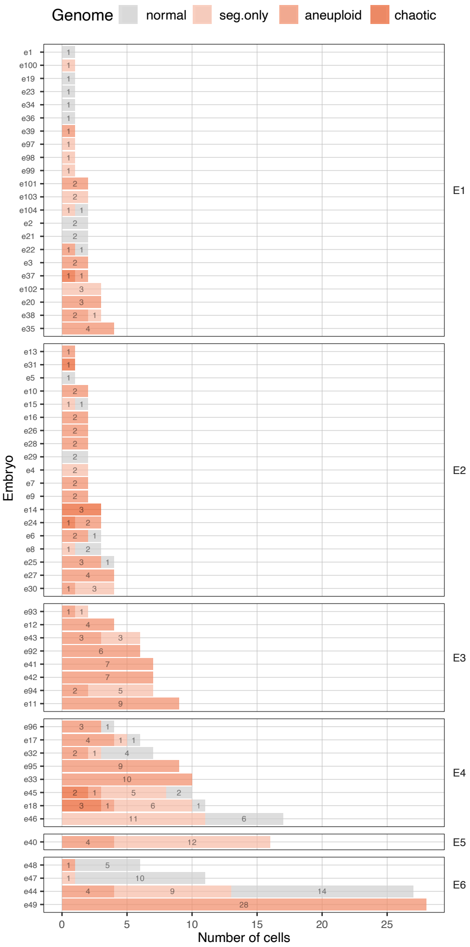

#### Extended Data Figure 2

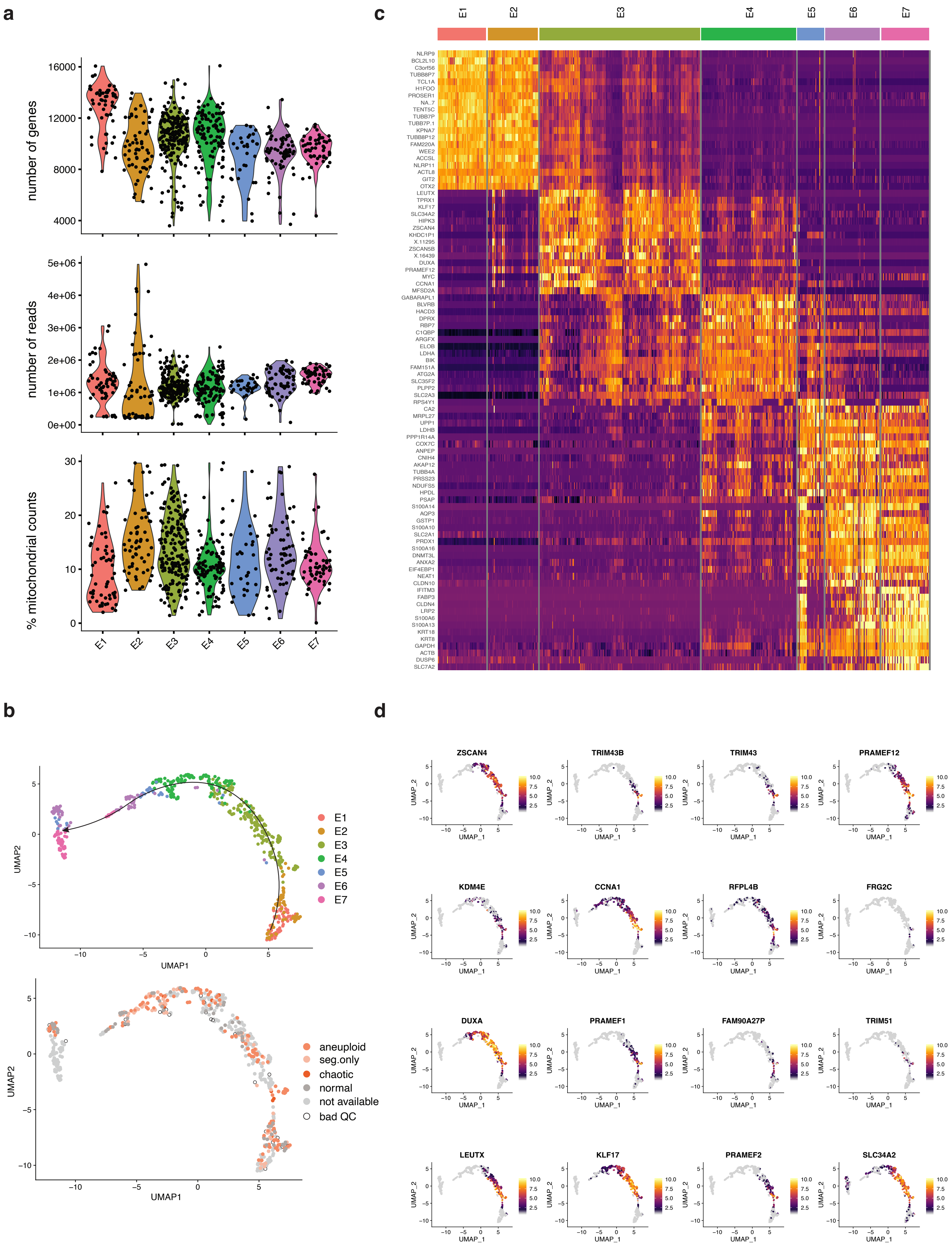

#### Extended Data Figure 3

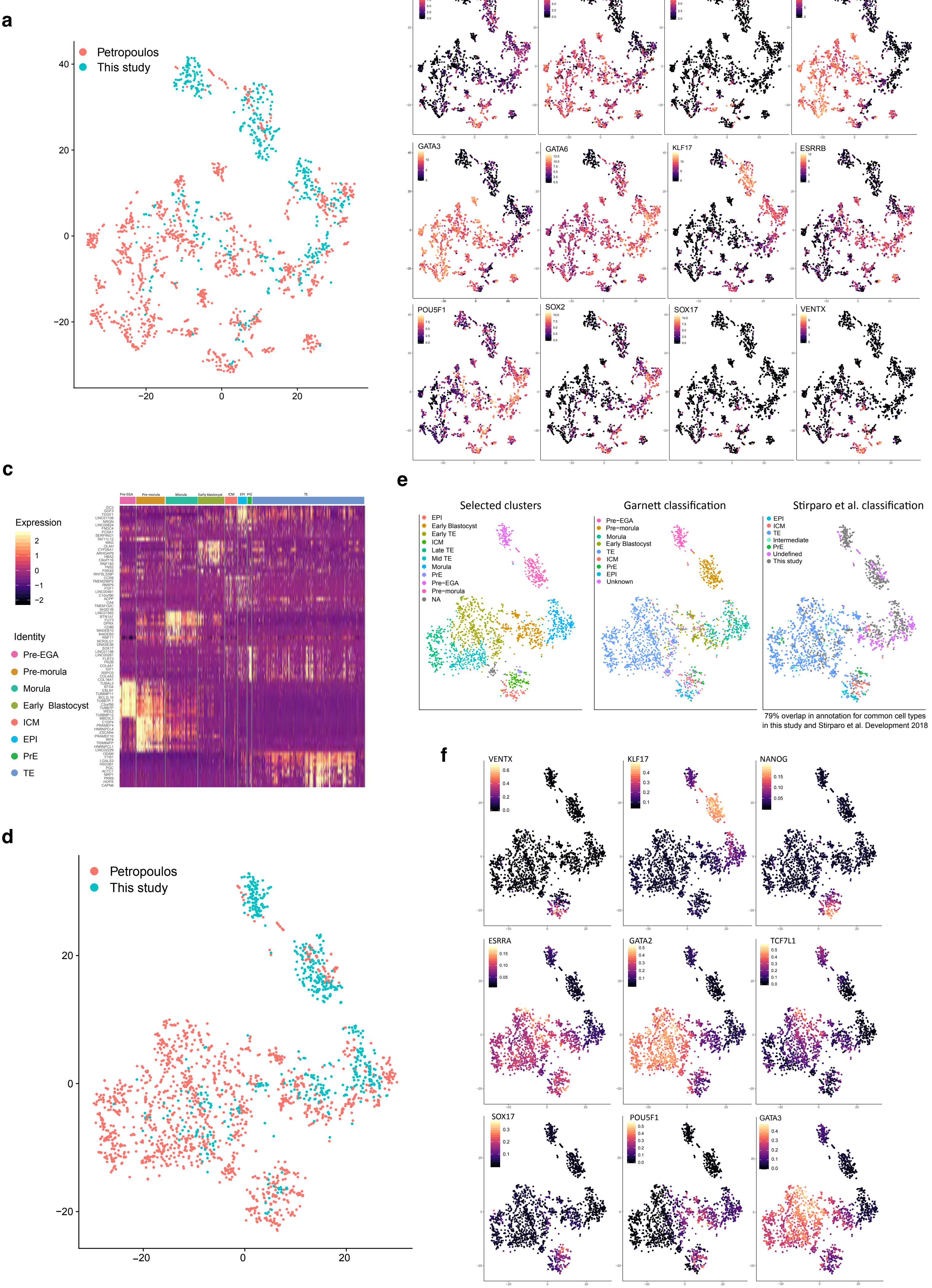

**Extended Data Figure 4**

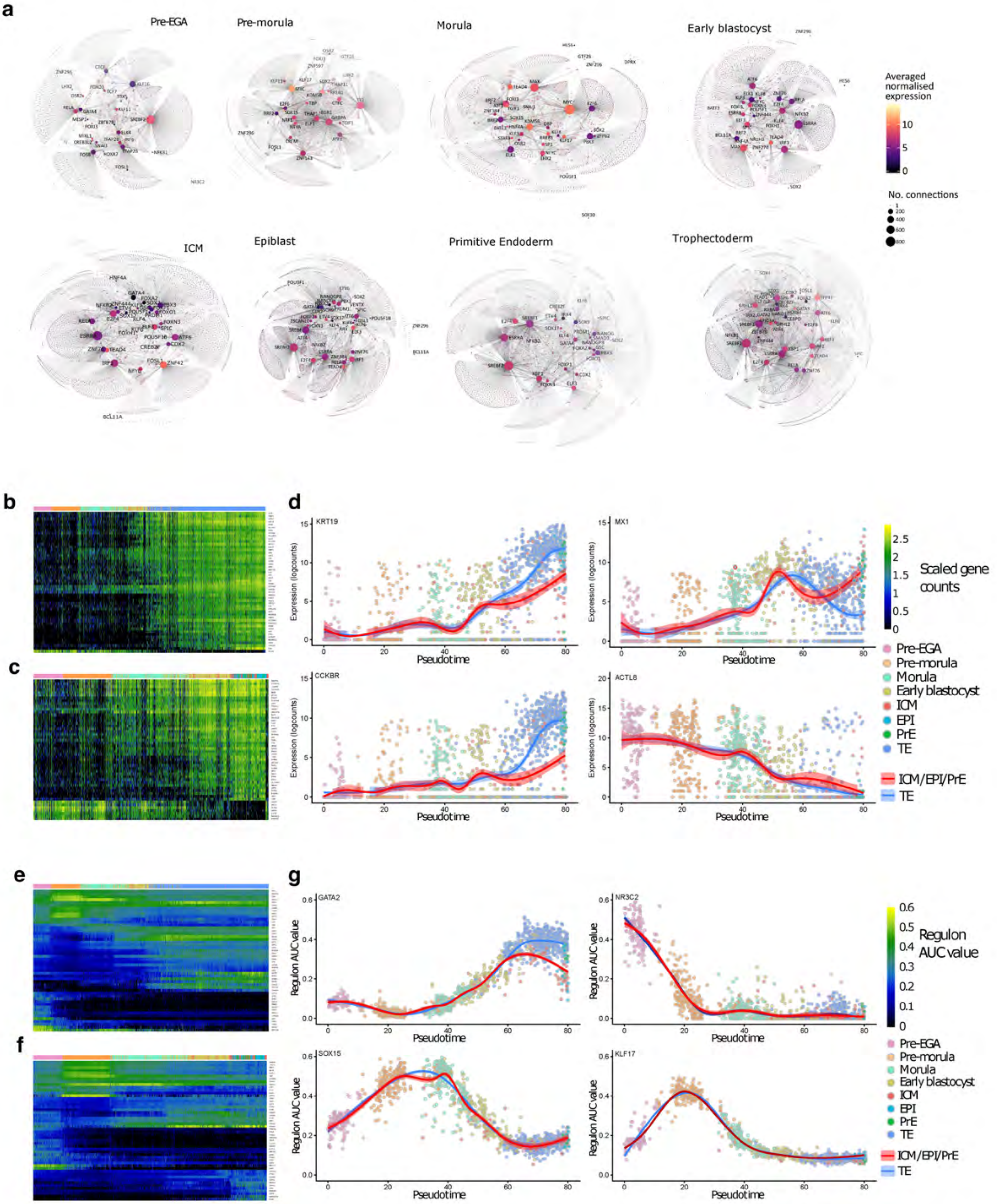
