## Extended Data Figures5-7 and Tables for "A multi-omics genome-and-transcriptome single-cell atlas of human preimplantation embryogenesis reveals the cellular and molecular impact of chromosome instability"

### Extended Data Figure 5

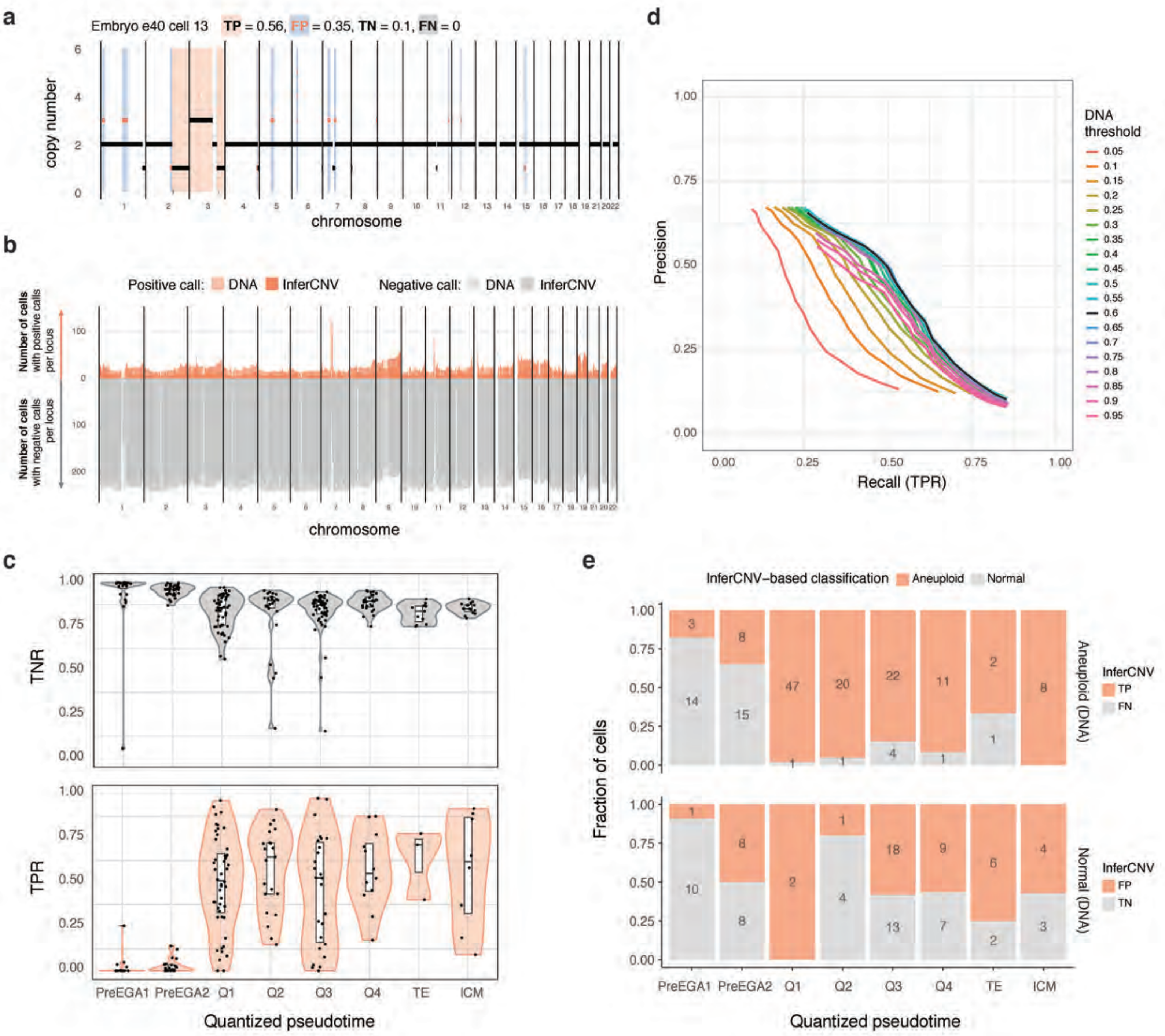

### Extended Data Figure 6

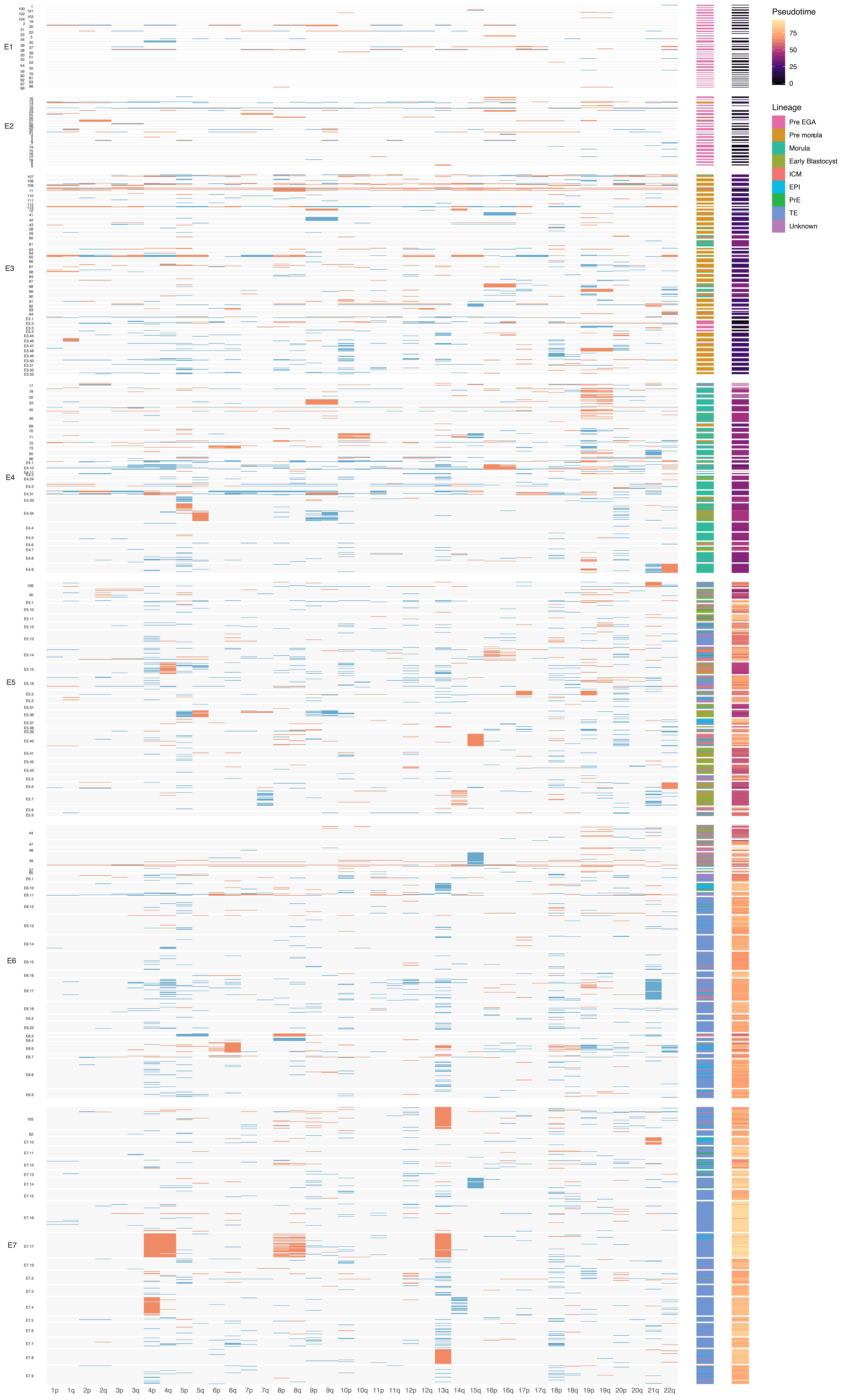

**Extended Data Figure 7**

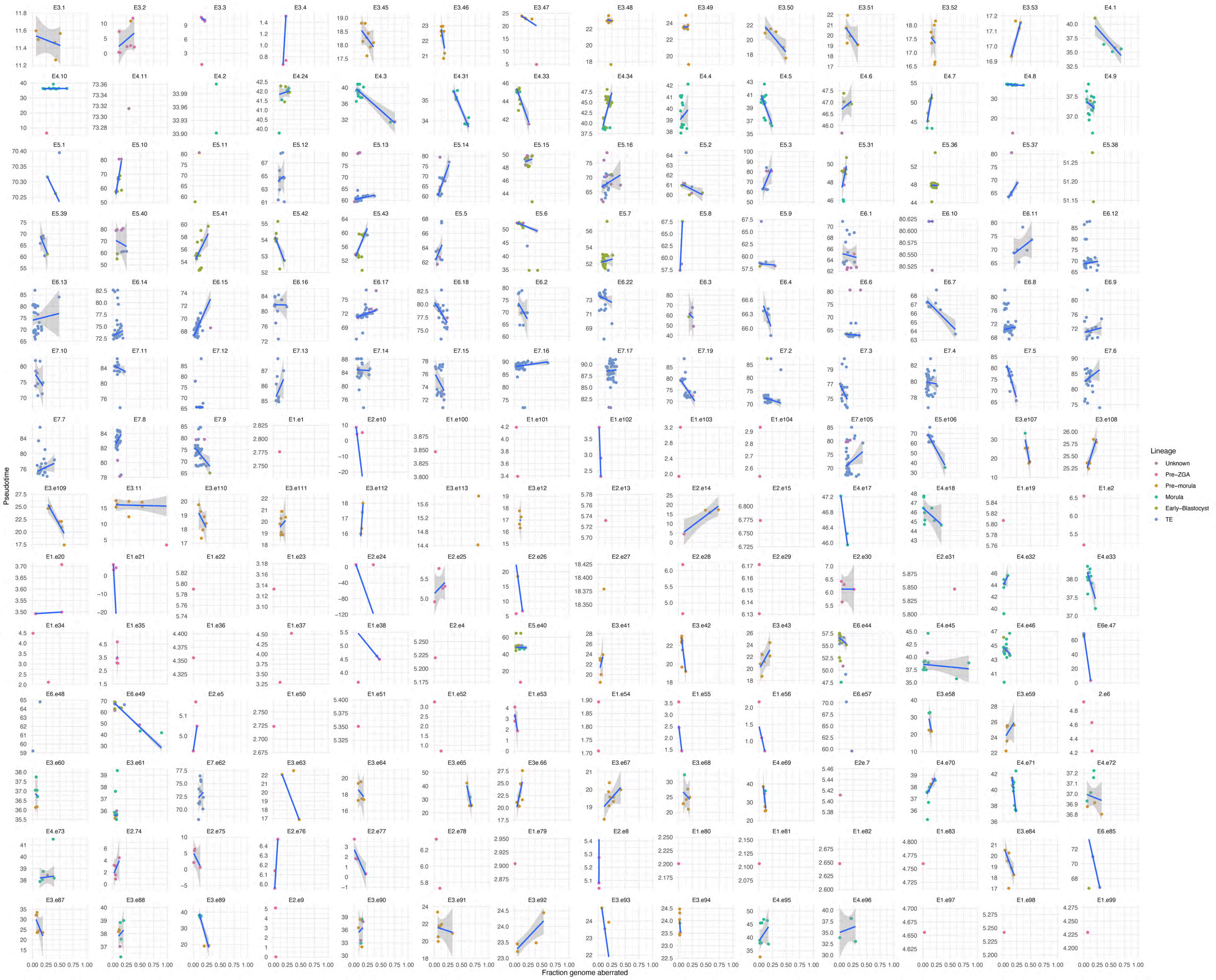

### EXTENDED DATA FIGURES

**Extended Data Fig. 1 Copy number variation from G&T-seq-derived DNA data. a.** Heatmap of raw genome-wide copy number profiles of 295 cells from 62 embryos ordered across embryonic days E1 to E6 and clustered by embryo. Gray represents normal euploid regions, shades of orange/red represent deletions and blue duplications, respectively. X chromosome copy number (CN) is normalised differently for male (X CN=1, grey) and female embryos (X CN=2, grey) and is left blank for embryos which sex could not be determined due to discordant X CN between cells. **b.** Genome-wide copy number and B-allele frequencies inferred by a Hidden-Markov model of 7 cells in embryo E3.e11. Monosomies are confirmed by the concurrent loss of heterozygosity observed in the same genomic regions. **c.** Number of cells per embryo coloured by type of genomic abnormality: normal, aneuploid, segmental only or chaotic when, respectively, none, a full chromosome, a chromosome segment or multiple segments in all chromosomes genome-wide were abnormal.

**Extended Data Fig. 2. RNA data from G&T-seq. a.** RNA-seq data quality parameters after G&T-seq in embryonic cells in each day of embryonic development E1-E6. **b.** Gene expression based UMAP plot of embryonic cells processed by G&T-seq coloured by day of development with a line showing the trajectory (top) or by genomic composition (bottom). **c.** Heatmap indicating relative expression levels of marker genes in each day of development. Gene functions are described in the main text. **d.** Gene expression based UMAP plots of embryonic cells processed by G&T-seq coloured by relative expression of genes related to embryonic genome activation described in De Iaco *et al.*, 2017[145].

**Extended Data Fig. 3. G&T-seq and SmartSeq2-integrated single-cell RNA dataset. a-b.** t-SNE generated from integrated gene counts from 2105 single cells, experimental origin and selected gene expression projected on top, respectively. **c.** Heat map showing the top 10 differentially expressed genes per lineage. **d.** t-SNE generated from regulon AUC values from 2105 single cells, experimental origin projected on top. **e.** Left, the regulon t-SNE with selected clusters. Middle and Right, lineage classification of cells using Garnett and lineage classification from Stirparo et al. (2018), respectively. **f.** The regulon t-SNE with selected regulon activity projected on top.

**Extended Data Fig. 4. Gene regulatory networks active in human preimplantation embryos. a.** Gene regulatory networks active within each lineage. Transcription factors are named on the network, other points are genes which exist within the regulon for said transcription factor. Size of circle depicts the number of genes (connections) that exist within each regulon for each transcription factor. Averaged normalised expression is projected on each gene. **b-c.** Heat map depicting the 50 most temporally expressed genes over the pseudotime trajectory towards the TE and ICM/EPI/PrE lineages, respectively. **d.** Selected genes showing expression over pseudotime, each point is a single cell. **e-f.** Heat map depicting the 50 most temporally expressed regulons over the pseudotime trajectory towards the TE and ICM/EPI/PrE lineages, respectively. **g.** Selected regulons showing activity over pseudotime, each point is a single cell.

**Extended Data Fig. 5. Benchmarking of genome-wide CN detection from RNA-seq data using inferCNV on cells with G&T-seq data. a.** Genome-wide DNA-seq and inferCNV-based copy number (CN) calls for one cell. True positive (TP), false positive (FP), true negative (TN) and false negative (FN) rates are calculated as a fraction of the genome and are coloured as indicated above the plot. **b.** Out of 254 cells across embryonic days E1-E6, number of cells with positive (CN≠2) and negative (CN=2) DNA-seq and inferCNV based-calls for each position in the genome. **c.** Distribution of inferCNV true positive and true negative rates (TPR and TNR) per cell in each pseudotime quantile. **d.** Precision and recall rates for all stages after EGA of different DNA- and inferCNV-based thresholds for aneuploid cell classification. Maximum AUC is achieved with a threshold of 0.6 fraction of abnormal chromosome arm. **e.** Fraction of DNA-based classified normal and aneuploid cells per pseudotime quantile coloured by the inferCNV-based classification.

**Extended Data Fig. 6. CN heatmap with lineage and pseudotime annotation of 2105 single cells in 196 embryos along preimplantation development.** DNA losses are depicted in red, while DNA gains in blue.

**Extended Data Fig. 7. Correlation between abnormal fraction of the genome and pseudotime.** Depicted for all analysed cells of each of the 192 embryos. The pseudotime corresponds to the TE branch. Embryos showing a significant negative correlation are displayed in **Fig. 4**.

### EXTENDED DATA TABLES

**Extended Data Table 1.** Number of embryos and cells with good quality scG&T-seq libraries.

| Dataset | Library type | | E1 | E2 | E3 | E4 | E5 | E6 | E7 | Total |
|---|---|---|---|---|---|---|---|---|---|---|
| G&T | RNA total | Cells | 58 | 60 | 192 | 113 | 31 | 65 | 57 | 576 |
| | | Embryos | 32 | 23 | 31 | 13 | 2 | 6 | 2 | 109 |
| | DNA total | Cells | 39 | 45 | 49 | 74 | 16 | 72 | 0 | 295 |
| | | Embryos | 22 | 19 | 8 | 8 | 1 | 4 | 0 | 62 |
| | DNA&RNA | Cells | 34 | 37 | 46 | 68 | 15 | 54 | 0 | 254 |
| | | Embryos | 19 | 18 | 8 | 8 | 1 | 4 | 0 | 58 |
| Petropoulos | RNA | Cells | 0 | 0 | 81 | 190 | 377 | 415 | 466 | 1529 |
| | | Embryos | 0 | 0 | 13 | 15 | 24 | 18 | 17 | 87 |
| Integrated | RNA | Cells | 58 | 60 | 273 | 303 | 408 | 480 | 523 | 2105 |
| | | Embryos | 32 | 23 | 44 | 28 | 26 | 24 | 19 | 196 |

**Extended Data Table 2.** Type and origin of genomic abnormalities detected by DNA-seq.

| | | | E1 | | E2 | | E3 | | E4 | | E5 | | E6 | | E7 | Total | |
|---|---|---|---|---|---|---|---|---|---|---|---|---|---|---|---|---|---|
| | | | n | % | n | % | n | % | n | % | n | % | n | % | n | n | % |
| | Total | Cells | 39 | | 45 | | 49 | | 74 | | 16 | | 72 | | 0 | 295 | |
| | | Embryos | 22 | | 19 | | 8 | | 8 | | 1 | | 4 | | 0 | 62 | |
| Origin | Normal | Cells | 11 | 28 | 8 | 18 | 0 | 0 | 15 | 20 | 0 | | 29 | 40 | 0 | 63 | 21 |
| | | Embryos | 7 | 32 | 2 | 11 | 0 | 0 | 0 | 0 | 0 | | 0 | 0 | 0 | 9 | 15 |
| | Mitotic | Cells | 25 | 64 | 35 | 78 | 36 | 73 | 63 | 85 | 16 | 100 | 43 | 60 | 0 | 218 | 74 |
| | | Embryos | 15 | 68 | 17 | 89 | 8 | 100 | 8 | 100 | 1 | 100 | 4 | 100 | 0 | 57 | 92 |
| | Meiotic (and likely) | Cells | 11 | 28 | 15 | 33 | 33 | 67 | 19 | 26 | 0 | | 28 | 39 | 0 | 106 | 36 |
| | | Embryos | 4 | 18 | 1 | 5 | 3 | 38 | 2 | 25 | 0 | | 1 | 25 | 0 | 11 | 18 |
| Type | Whole chromosome | Cells | 16 | 41 | 25 | 56 | 39 | 80 | 30 | 41 | 4 | 25 | 33 | 46 | 0 | 147 | 50 |
| | | Embryos | 8 | 36 | 12 | 63 | 8 | 100 | 7 | 88 | 1 | 100 | 3 | 75 | 0 | 39 | 63 |
| | Segmental only | Cells | 12 | 31 | 7 | 16 | 10 | 20 | 24 | 32 | 12 | 75 | 10 | 14 | 0 | 75 | 25 |
| | | Embryos | 9 | 41 | 4 | 21 | 3 | 38 | 5 | 63 | 1 | 100 | 2 | 50 | 0 | 24 | 39 |
| | Chaotic | Cells | 1 | 3 | 5 | 11 | 0 | | 5 | 7 | 0 | | 0 | | 0 | 11 | 4 |
| | | Embryos | 1 | 5 | 3 | 16 | 0 | | 2 | 25 | 0 | | 0 | | 0 | 6 | |

**Extended Data Table 3.** DNA-seq vs InferCNV based cell classification (>60% chr arm) per stage.

| | | E1 | | E2 | | E3 | | E4 | | E5 | | E6 | | E7 | | Total | |
|---|---|---|---|---|---|---|---|---|---|---|---|---|---|---|---|---|---|
| | | n | % | n | % | n | % | n | % | n | % | n | % | n | % | n | % |
| Total cells DNA&RNA | | 34 | | 37 | | 46 | | 68 | | 15 | | 54 | | 0 | | 254 | |
| DNA-seq | aneuploid | 16 | 47.1 | 28 | 76.0 | 44 | 95.7 | 38 | 55.9 | 8 | 53.3 | 24 | 44.4 | 0 | | 158 | 62.2 |
| InferCNV | aneuploid | 6 | 17.6 | 18 | 49.0 | 45 | 97.8 | 49 | 72.1 | 10 | 66.7 | 42 | 77.8 | 0 | | 170 | 66.9 |
| 1-TNR=* | false positives | 3 | 16.7 | 6 | 66.7 | 2 | 100 | 15 | 50.0 | 4 | 57.1 | 19 | 63.3 | 0 | | 49 | 51.0 |
| 1-TPR=** | false negatives | 13 | 81.3 | 16 | 57.1 | 1 | 2.3 | 4 | 10.5 | 2 | 25.0 | 1 | 4.2 | 0 | | 37 | 23.4 |

*inferCNV aneuploid/DNA normal; ** inferCNV normal/ DNA-seq aneuploid

**Extended Data Table 4.** DNA-seq vs InferCNV based cell classification (>60% chr arm) per pseudotime bin.

| | | PreEGA1 | | PreEGA2 | | Q1 | | Q2 | | Q3 | | Q4 | | ICM | | TE | | Total | |
|---|---|---|---|---|---|---|---|---|---|---|---|---|---|---|---|---|---|---|---|
| | | n | % | n | % | n | % | n | % | n | % | n | % | n | % | n | % | n | % |
| Total cells DNA&RNA | | 28 | | 39 | | 50 | | 26 | | 57 | | 28 | | 15 | | 11 | | 254 | |
| DNA-seq | aneuploid | 17 | 60.7 | 23 | 59.0 | 48 | 96.0 | 21 | 80.8 | 26 | 45.6 | 12 | 42.9 | 8 | 53.3 | 3 | 27.3 | 158 | 62.2 |
| InferCNV | aneuploid | 4 | 14.3 | 16 | 41.0 | 49 | 98.0 | 21 | 80.8 | 40 | 70.2 | 20 | 71.4 | 12 | 80.0 | 8 | 72.7 | 170 | 66.9 |
| 1-TNR=* | false positives | 1 | 9.1 | 8 | 50.0 | 2 | 100 | 1 | 20.0 | 18 | 58.1 | 9 | 56.3 | 4 | 57.1 | 6 | 75.0 | 49 | 51.0 |
| 1-TPR=** | false negatives | 14 | 82.3 | 15 | 65.2 | 1 | 2.1 | 1 | 4.8 | 4 | 15.4 | 1 | 8.3 | 0 | | 1 | 33.3 | 37 | 23.4 |

*inferCNV aneuploid/DNA normal; ** inferCNV normal/ DNA-seq aneuploid

**Extended Data Table 5.** Integrated Dataset (includes E1 and E2 embryos with only RNA) - classified according to inferCNV or by DNA-seq when available.

| | | E1 | | E2 | | E3 | | E4 | | E5 | | E6 | | E7 | | Tota l | |
|---|---|---|---|---|---|---|---|---|---|---|---|---|---|---|---|---|---|
| | | n | % | n | % | n | % | n | % | n | % | n | % | n | % | n | % |
| Total | Cells | 58 | | 60 | | 273 | | 303 | | 408 | | 480 | | 523 | | 2105 | |
| | Embryos | 32 | | 23 | | 44 | | 28 | | 26 | | 24 | | 19 | | 196 | |
| Aneuploid | Cells | 21 | 36.2 | 35 | 58.3 | 219 | 80.2 | 222 | 73.3 | 303 | 74.3 | 289 | 60.2 | 402 | 76.9 | 1491 | 70.8 |
| | Embryos | 12 | 38.0 | 19 | 83.0 | 44 | 100 | 28 | 100 | 26 | 100 | 24 | 100 | 19 | 100 | 172 | 88.0 |

**Extended Data Table 6.** Dosage sensitive genes for deletions and duplications classified by function.

| Function group | Del | Dup | Overlap | overlap genes | Other genes del | Other genes dup |
|---|---|---|---|---|---|---|
| Transcription, transcription regulation and RNA processing | 16 | 12 | 0 | - | *DHX15, HNRNPDL, EXOSC8, RBM47, GTF2A2, HNRNPD, SNU13, SNRPD3, POLR2K, POLR2F, ELOC, PLRG1, SF3B1, EWSR1, PRPF40A, MATR3, IPO5* | *POLR2L, SSU72, FUS, PRPF8, POP7, LRPPRC, U2AF1, STRAP, EDF1, HNRNPAB, SF3B2, ELOB* |
| Mitochondria related (respiratory chain, membrane, mitochondrial genome regulation) | 16 | 16 | 4 | *MRPS34, CYC1, AURKAIP1, ATAD3B* | *DNAJC15, ATP5ME, GRPEL1, NDUFC1, TMEM70, UQCR10, UQCRB, MRPL15, GRSF1, NDUFB9, TFB2M, MRPL13* | *COX8A, MTATP6P1, CHCHD2, COX4I1, ATP5F1E, NDUFB10, ATP5PO, NDUFS8, MRPL11, ATP5PF, COX5A, SLC25A39* |
| Ribosomal components and regulators | 15 | 12 | 7 | *RPL8, RPS2, RPLP2, RPL4, RPL13, FAU, RPLP1* | *RPL34, RPS20, POLR1D, ABCE1, RPL9, RPS14, LYAR, GTF3A* | *RPS21, RPL13, RPL3, NOP10, RPLP1, RRP7A, GTPBP4* |
| Chromatin remodeling, histones and histone modification | 11 | 10 | 5 | *H2AFZ, SAP18, WDR5, HMGB2, DNTTIP2* | *MRFAP1, SMARCA5, RBPJ, TET2, HDAC2, ENY2* | *HMGN1, NTMT1, IST1H4C, DNMT3L, WDR5, ZNHIT1* |
| Translation, post-translational modifications and protein transport | 10 | 11 | 3 | *EIF3E, NAA15, EIF2S1* | *TMEM33, EIF3H, SRP72, TMEM167A, EIF5A, TOMM22* | *CCT8, SNX2, EIF3B, LMAN2, RAB11A, TBC1D23, QARS, EMC4* |
| Protein degradation, autophagy and apoptosis | 9 | 9 | 2 | *UBE2D3, PSMA3* | *POMP, UBE2K, NEDD8, PSMB5, PSMA4, DCAF13, BCLAF1* | *OTUB1, ATG3, SIVA1, PSMC2, CTSD, ATG5, LAMTOR1* |
| Cell cycle regulation, cytoskeleton, DNA replication, proliferation | 8 | 8 | 2 | *MAD2L1, CFL1* | *PAICS, CCNA2, TPT1, CCNB1, NAA50, DRG1* | *PFN1, ANAPC11, MCM5, CCNC, CAPZA1, PTTG1IP* |
| Signal transduction | 5 | 11 | 3 | *YWHAZ, AZIN1, RPS19BP1* | *ANXA5, ANXA3* | *GNAS, PPP1CA, GNAI3, CALM1, MLF2, NRBP1, SPINT1, PTP4A1* |
| Molecule transport and membrane | 4 | 3 | 0 | - | - | - |

| | | | | | | |
|---|---|---|---|---|---|---|
| Metabolism | 3 | 5 | 1 | *ALDOA* | *FDFT1, AMD1* | *FASN, LDHB, CKB, DHCR7* |
| Oxidative stress | 1 | 3 | 1 | *TXN2* | - | *PRDX5, SOD1* |
| X inactivation | 1 | 0 | 0 | - | *ZFP42* | - |
| Stemness | 1 | 0 | 0 | - | *PROM1* | - |
| Total | 100 | 100 | 28 | | | |

**Extended Data Table 7.** Physical vs. molecular pseudotime age of euploid and aneuploid cells.

| Stage | P-value | Mean pseudotime aneuploid cells | Mean pseudotime euploid cells | Ratio |
|---|---|---|---|---|
| E3 | 0.0001266095056 | 21.75623578 | 26.53456626 | 0.8199205355 |
| E4 | 0.0001767149992 | 39.61980112 | 41.20499993 | 0.9615289695 |
| E5 | 0.0126514703 | 61.68913163 | 63.87375735 | 0.9657977578 |
| E6 | 0.3657318565 | 70.8525891 | 71.76576034 | 0.9872756697 |
| E7 | 0.02436889346 | 79.46281724 | 80.88128303 | 0.9824623728 |
| E1 | 0.0420284856 | 3.068237949 | 3.846479983 | 0.7976742275 |
| E2 | 0.774615953 | 6.428643317 | 4.817163948 | 1.334528653 |

**Extended Data Table 8.** Significant differentially expressed genes in aneuploid cells compared to euploid classified by function based on literature curation. One gene can be in multiple categories.

| Lineage group | In aneu-ploid cells | Genes (n) | Function category | Genes in category | Gene names and references |
|---|---|---|---|---|---|
| Post ZGA | Up | 5 | Oocyte related | 5 | *BPGM*[1], *CD59*[2], *GSKIP*[3,4], *SHOC2*[4,5], *ZP2*[6] |
| | Down | 22 | Nucleotide metabolism | 2 | *SHPK*[7], *UCK2*[8] |
| | | | mRNA stability and processing | 2 | *LRPPRC*[9], *PABPC4*[10] |
| | | | Ribosomal components | 5 | *RACK1*[11], *RPL3*, *RPL10A*, *RPL14*, *RPL22L1*, *RPL36AL* [12,13] |
| | | | Ribosomal binding protein | 1 | *RPS19BP1*[14] |
| | | | Translation elongation | 1 | *EEF1D*[15] |
| | | | Mitochondrial ribosomal proteins (MRPs) | 4 | *MRPS15*, *MRPL11*, *MRPL34*, *MRPS12*[16] |
| | | | Respiratory chain | 1 | *LRPPRC*[17,18], *NUBPL*[19] |
| | | | Proteasome | 1 | *PSMB5*[20] |
| | | | Linked to apoptosis upon downregulation and/or upregulated in cancer | 16 | *ARHGEF39*[21], *LRPPRC*[9], MRPs[22], *PABPC4*[23,24], *PSMB5*[20], *RACK1*[25,26], *RPL14*[27,28], *RPL10A*[29], *RPL22L1*[30–32], *RPS19BP1*[33], *TRIM28*[34], *UCK2*[35], *WWC1*[36,37] |
| | | | Essential for embryonic development | 9 | *LRPPRC*[38], MRPs[39], *PABPC4*[40] , *RPL10A*[29], *RPL22L1*[41], *TRIM28*[42] |
| ICM | Up | 1 | Signal transduction | 1 | *PTHP1*[43,44] |
| | Down | 2 | Ubiquitination | 1 | *UBE2J1*[45–47] |
| | | | Protein trafficking | 1 | *SNX3*[48] |
| TE | Up | 41 | Pro-proliferation | | |
| | | | Promote proliferation or survival | 9 | *CCND1*[49], *FAM9C*[50], *FGFR3*[51], *MIB2*[52], *PRPF19*[53], *PURPL*[54], *PVT1*[55], *SDF4*[56], *TMEM54*[57] |
| | | | Upregulated in cancer | 20 | *AOPEP*[58], *CAPG*[59], *CHI3L2*[60], *CST3*[61], *EPAS1*[62], *FAM9C*[50], *FBLN1*[63], *FGFR3*[51], *FTH1*[64], *IFITM3*[65], *NCOR2*[66], *NECTIN1*[67], *NQO1*[68],*PHPT1*[69], *PURPL*[54], *PVT1*[55], *SDF4*[70], *SFN*[71], *TMEM54*[57], *TNFRSF10B*[72] |
| | | | Anti-proliferation | | |
| | | | Induce cell cycle arrest or growth retardation | 7 | *CDKN1A*[73], *FDXR*[74], *NQO1*[68], *RPS27L*[75], *PRPF19*[76], *SESN1*[77], *SFN*[71] |
| | | | Trigger apoptosis signalling | 3 | *EDA2R*[78], *NQO1*[68], *RPS27L*[79], *TNFRSF10B*[80] |
| | | | Targets of p53 | 8 | *CDKN1A*[73], *EDA2R*[81], *FTH1*[82], *FDXR*[74], *PHLDA3*[83], *RPS27L*[75], *SESN1*[77], *SFN*[71] |
| | | | Response to DNA damage, oxidative stress, radiation | 14 | *CCND1*[84], *CDKN1A*[85,86], *EPAS1*[87], *FDXR*[88], *FTH1*[82], *NQO1*[68], *PHLDA3*[89], *PHPT1*[88], *PRPF19*[76], *RPS27L*[90], *SESN1*[88], *SFN*[91], *TMEM54*[92], *TNFRSF10B*[84] |
| | | | Implantation related | | |
| | | | Implantation / expressed in trophoblasts | 14 | *CAPG*[59], *CDKN1A*[93], *COLT1*[94], *CST3*[61], *EPAS1*[95], *F3*[96], *FBLN1*[97], *FGFR3*[98], *FTH1*[64], *IFITM3*[99], *KRT7*[100,101], *NECTIN1*[102], *PVT1*[103], *TNFRSF10B*[104] |
| | | | Iron transport and metabolism | 3 | *FTL*[105], *FTH1*[82], *FDXR*[74] |
| | | | WNT signalling | 3 | *CCND1*[106], *IFITM3*[65], *LRP1*[107] |
| | | | Other | | |
| | | | Pluripotency regulation | 1 | *TENT5B*[108] |
| | | | Cytoskeleton and its regulation | 2 | *BBS9*[109], *PLEKHH3*[110] |
| | | | Suppressor of cytokine signalling | 1 | *ASB1*[111] |
| | | | Ribosomal components | 2 | *RPS19*, *RPS27L*[12,13] |

| Lineage group | In aneu-ploid cells | Genes (n) | Function category | Genes in category | Gene names and references |
|---|---|---|---|---|---|
| TE | Down | 90 | Pro-proliferation | | |
| | | | Proliferation | 18 | *AIF1L*[112], *AK4*[113], *CBS*[114], *DANCR*[115], *ENOSF1*[116], *IMPDH2*[117], *LETMD1*[118], *MOB1B*[119], *MTHFD2*[120], *OGDH*[121], *PAICS* [122], *PFN2*[123], *PHGDH*[124], *PLOD1*[125], *RBM14*[126], *STC2*[127], *UAP1L1*[128], *ZNF165*[129] |
| | | | Upregulated in cancer | 29 | *ACSL4*[130], *ACSL6*[131], *AK4*[113], *AMD1*[132], *ANXA5*[133], *ASNS*[134], *CHAC1*[135], *CLDN10*[136], *DANCR*[115], *DDIT4*[137], *ENOPH1*[138], *HNRNPD*[139], *HK2*[140], *IMPDH2*[117], *LETMD1*[141], *LDHA*[142],*LDHB*[142], *MTHFD2*[120],*OGDH*[143], *P4HA1*[144], *PAICS* [122], *PFN2*[145], *PHGDH*[124], *PLOD1*[125], *PSAT1*[146], *PYCR1*[147], *RCC1*[148], *RPL22L1*[32] , *STC2*[127], *TARS*[149], *UAP1L1*[128], *ZNF165*[129] |
| | | | p53 inhibitors/inhibited by p53 | 2 | *ASNS*[134], *DHX15*[150] |
| | | | Context dependent pro and anti proliferation | 2 | *DDIT4*[151], *DHX15*[152] |
| | | | Anti-proliferation | | |
| | | | Apoptosis | 6 | *CHAC1*[135], *DDIT4*[151], *DNAJA3*[153], *MRFAP1*[154], *RBM47*[155], *TRIM13*[156] |
| | | | Downregulated in cancer | 4 | *ACSL6*[131], *ANXA5*[133], *MRFAP1*[157], *SLC25A43*[158], *RBM47*[155], *RCC1*[148] |
| | | | p53 targets/activators | 7 | *DDIT4*[159], *DNAJA3*[153], *GTF3A*[160] , *MAP4*[161], *RBM47*[155], *TRIM13*[156],*WARS*[162] |
| | | | Stress response | | |
| | | | ER stress/UPR/Autophagy | 9 | *ACSL4*[163], *CHAC1*[135], *CHMP4B*[164], *DDIT4*[151], *DHX15*[165], *HK2*[140], *LDHB*[142], *STC2*[127], *TRIM13*[166] |
| | | | Hypoxia/Oxidative stress | 6 | *AK4*[167], *CHAC1*[135], *DDIT4*[168], *LDHA*[169], *PYCR1*[170], *STC2*[127] |
| | | | DNA damage response | 3 | *RBM14*[171], *PAICS*[172], *DDIT4*[151] |
| | | | Protein degradation | 2 | *CHMP4B*[173], *TRIM13*[47] |
| | | | Immune response | 4 | *IL34*[174], *RGS13*[175], *TRIM13*[176], *WARS*[162] |
| | | | Cell cycle regulation | | |
| | | | DNA replication | 2 | *MCM3AP*[177], *MTHFD2*[120] |
| | | | Mitosis regulation | 11 | *CHMP4B*[178], *KIF21A*[179], *MAP4*[180], *MRFAP1*[154], *NUP58*[181,182], *PFN2*[183], *RBM14*[184], *RCC1*[148], *SLC25A43*[185] |
| | | | Cytoskeleton and its regulation | 9 | *AIF1L*[186], *ARHGAP18*[187], *KIF21A*[179], *KLHL23*[188], *MAP4*[180], *MOB1B*[189], *NUP58*[181,182], *PFN2*[183], *RBM14*[184] |
| | | | Nucleotide synthesis/homeostasis | 4 | *AK4*[190], *IMPDH2*[117], *MTHFD2*[191], *PAICS*[192] |
| | | | Transcripton and its regulation | | |
| | | | Transcription | 5 | *AFF1*[193], *GTF3A*[160], *IMPDH2*[117], *MED18*[194], *MTHFD2*[120], *NPM3*[195], *ZNF165*[129] |
| | | | mRNA splicing | 3 | *DHX15*[150], *MBNL3*[196], *RBM47*[155] |
| | | | Chromatin remodelling | 5 | *MRFAP1*[154], *NPM3*[195], *OGDH*[143], *RBBP4*[197], *RCC1*[148] |
| | | | Pluripotency regulation | 9 | *AMD1*[198], *MBNL3*[199], *MTHFD2*[120], *NPM3*[200], *PSAT1*[201], *RBBP4*[202], *RBM14*[203], *RBM47*[155], *SLC38A2*[204] |
| | | | Translation | | |
| | | | Translation and its regulation | 4 | *ATXN2L*[205], *HNRNPD*[206], *IMPDH2*[117], *NUFIP2*[207] |
| | | | Aminoacyl-tRNA synthetases | 5 | *EPRS, IARS, TARS, WARS, YARS*[208,209] |

| | | | | | |
|---|---|---|---|---|---|
| | | | Ribosome regulation/biogenensis | 4 | *DHX15*[210], *GTF3A*[160], *NPM3*[211], *RPL22L1*[212] |

(Continues in next page)

(Continues from previous page)

| Lineage group | In aneu-ploid cells | Genes (n) | Function category | Genes in category | Gene names and references |
|---|---|---|---|---|---|
| TE | Down | 90 | Metabolism | | |
| | | | Aminoacid metabolism | | *ACSL4*[213], *AMD1*[214], *ASNS*[134], *CBS*[215], *CHAC1*[135], *ENOPH1*[138], *IVD*[216], *MTR*[217], *PHGDH*[124], *PSAT1*[218], *PYCR1*[147] |
| | | | Carbohydrate metabolism | | *AGL*[219], *ENOSF1*[220], *DDIT4*[151], *HK2*[140], *LDHB*[142], *LDHA*[142], *OGDH*[143], *SLC39A14*[221] |
| | | | Lipid metabolism | | *ACSL6*[131], *ACSL4*[131], *DDIT4*[151], *EPRS*[222], *RNF145*[223] |
| | | | Mitochondrial | | *AK4*[224], *DNAJA3*[153], *ENOSF1*[220], *HK2*[140], *IVD*[216], *LETMD1*[118], *MIPEP*[225], *MTHFD2*[191], *OGDH*[143], *SLC25A43*[185] |
| | | | Cell communication/molecular transport | | |
| | | | Cell communication | | *ADGRE5*[226], *ANXA5*[227], *CLDN10*[228], *CPN3*[229], *STXBP6*[230], *RGS13*[231], *STC2*[127], *TARS*[208], *WARS*[208], *YARS*[208] |
| | | | Membrane trafficking | | *ANXA5*[232], *CHMP4B*[173], *ERGIC1*[233], *STXBP6*[230], *TRIM13*[234] |
| | | | Extracellular matrix | | *P4HA1*[235], *PLOD1*[125], *PHGDH*[236], *PYCR1*[236] |
| | | | Molecule transport | | *ABCC4*[237], *NUP58*[238], *SLC15A2*[239], *RCC1*[240], *IPO5*[241], *SLC39A14*[242], *SLC38A2*[243] |
| | | | Signal transduction | | |
| | | | Glutathione | | *CHAC1*[135] |
| | | | Phosphate homeostasis | | *STC2*[127], *XPR1*[244] |
| | | | Calcium and inositol signalling | | *CPNE3*[245], *PLCG1*[246], *PFN2*[145], *STC2*[127] |
| | | | WNT signalling | | *ABCC4*[247], *OGDH*[121], *PLOD1*[248], *RBM14* |
| | | | mTORC | | *ADGRE5*, *CBS*[114], *DDIT4*[151], *ENOPH1*[138], *EPRS*[222], *HK2*[140], *IMPDH2*[249], *LDHB*[142] |
| | | | Hippo-YAP/TAZ | | *ARHGAP18*[187], *MOB1B*[250], *PLOD1*[251] |
| | | | NF-κß | | *DNAJA3*[252], *TRIM13*[176] |
| | | | Inhibitor of TGF-ß | | *PFN2*[145] |
| | | | Other signal transduction pathways | | *CPNE3*[253], *IL34*[174], *PLOD1*[125], *RGS13*[231], *EPRS, IARS, TARS, WARS, YARS*[208] |
| | | | Implantation related | | |
| | | | Implantation / expressed in trophoblasts | | *ABCC4*[247], *ACP5*[254], *ADGRE5*[255], *ANXA5*[232], *CBS*[114], *CLDN10*[256], *FUT1*[257], *P4HA1*[258], *IL34*[259], *LDHB*[260], *MTR*[217], *SLC38A2*[243], *STC2*[261] |
| | | | EMT | | *IMPDH2*[249], *KLHL23*[188] (inhibitor), *NUP58*[262], *PFN2*[123], *RBM47*[263], *RPL22L1*[212] |
| | | | Iron homeostasis | | *ACP5*[264], *MIPEP*[225], *SLC39A14*[265] |
| | | | Angiogenesis regulation | | *WARS, YARS*[266] |

8. Suzuki, N. N. *et al.* (2004) *Struct. Lond. Engl. 1993* 12, 751–764.
9. Cui, J. *et al.* (2019) *Front. Physiol.* 10, 595.
10. Jalkanen, A. L. *et al.* (2014) *Semin. Cell Dev. Biol.* 34, 24–32.
11. Johnson, A. G. *et al.* (2019) *RNA* 25, 881–895.
12. Wilson, D. N. & Cate, J. H. D. (2012) *Cold Spring Harb. Perspect. Biol.* 4, a011536.
13. Zhou, X. *et al.* (2015) *J. Mol. Cell Biol.* 7, 92–104.
14. Knight, J. R. P. *et al.* (2013) *Nucleic Acids Res.* 41, 4185–4197.
15. Gingras, A.-C. *et al.* (1999) *Annu. Rev. Biochem.* 68, 913–963.
16. Cheong, A. *et al.* (2020) *Gene Expr. Patterns GEP* 38, 119147.
17. F, X. *et al.* (2012) *Biochem. J.* 441,.
18. Aibara, S. *et al.* (2020) *eLife* 9, e58362.
19. Sheftel, A. D. *et al.* (2009) *Mol. Cell. Biol.* 29, 6059–6073.
20. Wei, W. *et al.* (2018) *Int. J. Biol. Markers* 33, 102–108.
21. Wang, S. *et al.* (2020) *Genet. Mol. Biol.* 43, e20190383.
22. Huang, G. *et al.* (2020) *Int. J. Mol. Sci.* 21, E8879.
23. Liu, D. *et al.* (2012) *J. Genet. Genomics Yi Chuan Xue Bao* 39, 369–374.
24. Jiang, X. *et al.* (2021) *Cancer Lett.* 499, 109–121.
25. Duff, D. & Long, A. (2017) *Cell. Signal.* 35, 250–255.
26. Xiao, T. *et al.* (2018) *Cell Death Dis.* 9, 1148.
27. Feng, Y. *et al.* (2019) *Biochem. Biophys. Res. Commun.* 513, 56–63.
28. Yu, C. *et al.* (2019) *Mol. Med. Rep.* 19, 2065–2076.
29. Palasin, K. *et al.* (2019) *Sci. Rep.* 9, 18130.
30. Rao, S. *et al.* (2019) *PloS One* 14, e0222392.
31. Liang, Z. *et al.* (2019) *Med. Oncol. Northwood Lond. Engl.* 36, 56.
32. Wu, N. *et al.* (2015) *PloS One* 10, e0143659.
33. Brooks, C. L. & Gu, W. (2009) *Nat. Rev. Cancer* 9, 123–128.
34. Czerwińska, P. *et al.* (2017) *J. Biomed. Sci.* 24, 63.
35. Yu, S. *et al.* (2019) *IUBMB Life* 71, 105–112.
36. Song, L. *et al.* (2019) *Front. Cell. Neurosci.* 13, 137.
37. Wilson, K. E. *et al.* (2016) *Genes* 7, E23.
38. Xu, F. *et al.* (2012) *Biochem. J.* 441, 275–283.
39. Cheong, A. *et al.* (2020) *Dev. Camb. Engl.* 147, dev188714.
40. Gorgoni, B. *et al.* (2011) *Proc. Natl. Acad. Sci. U. S. A.* 108, 7844–7849.
41. Zhang, Y. *et al.* (2017) *Cell Rep.* 18, 545–556.
42. Messerschmidt, D. M. *et al.* (2012) *Science* 335, 1499–1502.
43. Attwood, P. V. (2013) *Biochim. Biophys. Acta* 1834, 470–478.
44. Fuhs, S. R. & Hunter, T. (2017) *Curr. Opin. Cell Biol.* 45, 8–16.
45. Lenk, U. *et al.* (2002) *J. Cell Sci.* 115, 3007–3014.
46. Elangovan, M. *et al.* (2017) *J. Cell Commun. Signal.* 11, 265–273.
47. Lopata, A. *et al.* (2020) *Int. J. Mol. Sci.* 21, E5369.
48. Harterink, M. *et al.* (2011) *Nat. Cell Biol.* 13, 914–923.
49. Qie, S. & Diehl, J. A. (2016) *J. Mol. Med. Berl. Ger.* 94, 1313–1326.
50. Zhou, J.-D. *et al.* (2013) *Oncol. Rep.* 30, 1275–1284.
51. Turner, N. & Grose, R. (2010) *Nat. Rev. Cancer* 10, 116–129.
52. Feltham, R. *et al.* (2018) *Cell Rep.* 23, 470–484.
53. Fortschegger, K. *et al.* (2007) *Mol. Cell. Biol.* 27, 3123–3130.
54. Li, X. L. *et al.* (2017) *Cell Rep.* 20, 2408–2423.
55. Wang, W. *et al.* (2019) *Front. Oncol.* 9, 609.
56. Chen, L. *et al.* (2014) *Cell Death Dis.* 5, e1219.
57. Kong, Y. *et al.* (2009) *Cell Cycle Georget. Tex* 8, 3552–3561.
58. Ding, L. *et al.* (2017) *Tumour Biol. J. Int. Soc. Oncodevelopmental Biol. Med.* 39, 1010428317691657.
59. Lang, Z. *et al.* (2019) *Pathol. Res. Pract.* 215, 152683.
60. Zhao, T. *et al.* (2020) *Signal Transduct. Target. Ther.* 5, 201.
61. Jiang, Y. *et al.* (2020) *Hematol. Amst. Neth.* 25, 457–463.
62. Tirpe, A. A. *et al.* (2019) *Int. J. Mol. Sci.* 20, E6140.
63. Obaya, A. J. *et al.* (2012) *Cancer Lett.* 325, 132–138.
64. Moroz, C. *et al.* (2002) *J. Biol. Chem.* 277, 12901–12905.
65. Rajapaksa, U. S. *et al.* (2020) *Front. Oncol.* 10, 593245.
66. Battaglia, S. *et al.* (2010) *Int. J. Cancer* 126, 2511–2519.
67. Fujiwara, T. *et al.* (Springer Japan, 2016). in *The Cadherin Superfamily: Key Regulators of Animal Development and Physiology* (eds. Suzuki, S. T. & Hirano, S.) 115–156.
68. Zhang, K. *et al.* (2018) *J. Med. Chem.* 61, 6983–7003.
69. Snezhkina, A. V. *et al.* (2016) *BMC Genomics* 17, 1011.
70. Gong, W. *et al.* (2021) *Int. J. Mol. Med.* 47, 435–443.
71. Huang, Y. *et al.* (2020) *Clin. Chim. Acta* 511, 50–58.
72. von Karstedt, S. *et al.* (2015) *Cancer Cell* 27, 561–573.
73. Kreis, N.-N. *et al.* (2019) *Cancers* 11, E1220.
74. Zhang, Y. *et al.* (2017) *Genes Dev.* 31, 1243–1256.
75. Xiong, X. *et al.* (2011) *Oncogene* 30, 1798–1811.
76. Mahajan, K. (2016) *Oncogene* 35, 2279–2286.
77. Rhee, S. G. & Bae, S. H. (2015) *Free Radic. Biol. Med.* 88, 205–211.
78. Sinha, S. K. & Chaudhary, P. M. (2004) *J. Biol. Chem.* 279, 41873–41881.
79. He, H. & Sun, Y. (2007) *Oncogene* 26, 2707–2716.
80. Shirley, S. *et al.* (2011) *Recent Patents Anticancer Drug Discov.* 6, 311–323.
81. Brosh, R. *et al.* (2010) *FEBS Lett.* 584, 2473–2477.
82. Funauchi, Y. *et al.* (2015) *Sci. Rep.* 5, 16497.
83. Kawase, T. *et al.* (2009) *Cell* 136, 535–550.
84. Chaudhry, M. A. (2008) *J. Biomed. Sci.* 15, 557–563.
85. Adiga, S. K. *et al.* (2007) *Oncogene* 26, 6141–6149.
86. Ock, S.-A. *et al.* (2020) *Cell Stress Chaperones* 25, 503–508.
87. Majmundar, A. J. *et al.* (2010) *Mol. Cell* 40, 294–309.
88. Paul, S. & Amundson, S. A. (2008) *Int. J. Radiat. Oncol. Biol. Phys.* 71, 1236–1244.
89. Leszczynska, K. B. *et al.* (2015) *J. Clin. Invest.* 125, 2385–2398.
90. Li, J. *et al.* (2007) *Cancer Res.* 67, 11317–11326.
91. Mackintosh, C. (2004) *Biochem. J.* 381, 329–342.
92. Kigoshi, Y. *et al.* (2015) *Sci. Rep.* 5, 12857.
93. Higuchi, S. *et al.* (2019) *Placenta* 85, 56–62.
94. Provost, P. *et al.* (2001) *Biochem. J.* 359, 255–263.
95. Sasagawa, T. *et al.* (2018) *Sci. Rep.* 8, 17375.
96. Girardi, G. (2011) *J. Thromb. Haemost. JTH* 9, 250–256.
97. Okada, H. *et al.* (2014) *J. Obstet. Gynaecol. Res.* 40, 1180–1187.
98. Ozawa, M. *et al.* (2013) *Reprod. Camb. Engl.* 145, 191–201.
99. Zani, A. *et al.* (2019) *J. Biol. Chem.* 294, 19844–19851.
100. Lim, H. Y. G. *et al.* (2020) *Nature* 585, 404–409.
101. Marchand, M. *et al.* (2011) *Biol. Reprod.* 84, 1258–1271.
102. Cuman, C. *et al.* (2015) *EBioMedicine* 2, 1528–1535.
103. Qin, X. *et al.* (2019) *Am. J. Reprod. Immunol. N. Y. N 1989* 81, e13048.
104. Fluhr, H. *et al.* (2013) *Reprod. Biol.* 13, 113–121.
105. Arosio, P. *et al.* (2015) *Biochem. J.* 472, 1–15.
106. Katoh, M. & Katoh, M. (2007) *Clin. Cancer Res. Off. J. Am. Assoc. Cancer Res.* 13, 4042–4045.
107. Terrand, J. *et al.* (2009) *J. Biol. Chem.* 284, 381–388.
108. Hu, J.-L. *et al.* (2020) *Nucleic Acids Res.* 48, 2733–2748.
109. Klink, B. U. *et al.* (2020) *eLife* 9, e53910.
110. Lemmon, M. A. *et al.* (2002) *FEBS Lett.* 513, 71–76.
111. Starr, R. *et al.* (1997) *Nature* 387, 917–921.
112. Zhang, Y. *et al.* (2016) *Cancer Res.* 76, 4535–4545.
113. Xin, F. *et al.* (2019) *Clin. Exp. Med.* 19, 525–534.
114. Cai, S. *et al.* (2021) *Cell Prolif.* 54, e12950.
115. Jin, S.-J. *et al.* (2019) *Front. Oncol.* 9, 1225.
116. Finckbeiner, S. *et al.* (2011) *Cell Biosci.* 1, 32.
117. Kofuji, S. & Sasaki, A. T. (2020) *J. Biochem. (Tokyo)* 168, 319–328.
118. Cho, G.-W. *et al.* (2007) *BMC Cell Biol.* 8, 50.
119. Praskova, M. *et al.* (2008) *Curr. Biol. CB* 18, 311–321.
120. Zhu, Z. & Leung, G. K. K. (2020) *Front. Oncol.* 10, 658.
121. Lu, X. *et al.* (2019) *OncoTargets Ther.* 12, 7489–7500.
122. Agarwal, S. *et al.* (2020) *Cancers* 12, E772.
123. Tang, Y.-N. *et al.* (2015) *Nat. Commun.* 6, 8230.
124. Zhao, X. *et al.* (2020) *Int. J. Biol. Sci.* 16, 1495–1506.
125. Qi, Y. & Xu, R. (2018) *Front. Cell Dev. Biol.* 6, 66.
126. Carbon, S. & Mungall, C. (2018).
127. Joshi, A. D. (2020) *Front. Endocrinol.* 11, 172.
128. Xiao, X. *et al.* (2021) *Mol. Carcinog.* 60, 179–187.
129. Gibbs, Z. A. *et al.* (2020) *eLife* 9, e57679.
130. Tang, Y. *et al.* (2018) *Oncol. Lett.* 16, 1390–1396.
131. Rossi Sebastiano, M. & Konstantinidou, G. (2019) *Int. J. Mol. Sci.* 20, E3624.
132. Zabala-Letona, A. *et al.* (2017) *Nature* 547, 109–113.
133. Peng, B. *et al.* (2014) *Clin. Chim. Acta Int. J. Clin. Chem.* 427, 42–48.
134. Chiu, M. *et al.* (2019) *Front. Oncol.* 9, 1480.
135. Bachhawat, A. K. & Kaur, A. (2017) *Antioxid. Redox Signal.* 27, 1200–1216.
136. Zhang, X. *et al.* (2019) *J. Cell Commun. Signal.* 13, 395–405.
137. Tirado-Hurtado, I. *et al.* (2018) *Front. Oncol.* 8, 106.
138. Wang, B. *et al.* (2021) *J. Cell. Physiol.* 236, 1184–1194.
139. Zucconi, B. E. & Wilson, G. M. (2011) *Front. Biosci. Landmark Ed.* 16, 2307–2325.
140. Roberts, D. J. & Miyamoto, S. (2015) *Cell Death Differ.* 22, 248–257.
141. Jung, S. S. *et al.* (2005) *Clin. Cancer Res. Off. J. Am. Assoc. Cancer Res.* 11, 7700–7708.
142. Urbańska, K. & Orzechowski, A. (2019) *Int. J. Mol. Sci.* 20, E2085.
143. Vatrinet, R. *et al.* (2017) *Cancer Metab.* 5, 3.
144. Eriksson, J. *et al.* (2020) *Mol. Oncol.* 14, 742–762.
145. Pimm, M. L. *et al.* (2020) *Int. Rev. Cell Mol. Biol.* 355, 155–204.
146. Wei, Z. *et al.* (2020) *Front. Cell Dev. Biol.* 8, 603837.
147. D'Aniello, C. *et al.* (2020) *Front. Oncol.* 10, 776.
148. Ren, X. *et al.* (2020) *Front. Mol. Biosci.* 7,.
149. Wellman, T. L. *et al.* (2014) *BMC Cancer* 14, 620.
150. McElderry, J. *et al.* (2019) *Hum. Mol. Genet.* 28, 4173–4185.
151. Britto, F. A. *et al.* (2020) *Am. J. Physiol. Cell Physiol.* 319, C807–C824.

152. Inesta-Vaquera, F. *et al.* (2018) *Life Sci. Alliance* 1, e201800092.
153. Ahn, B. Y. *et al.* (2010) *Oncogene* 29, 1155–1166.
154. Li, D.-Z. *et al.* (2017) *Oncotarget* 8, 97178–97186.
155. Shivalingappa, P. K. M. *et al.* (2021) *Mol. Cell. Biochem.* 476, 4493–4505.
156. Hatakeyama, S. (2011) *Nat. Rev. Cancer* 11, 792–804.
157. Hu, L. *et al.* (2018) *Eur. Rev. Med. Pharmacol. Sci.* 22, 8273–8280.
158. Tina, E. *et al.* (2012) *BMC Cancer* 12, 350.
159. Ellisen, L. W. *et al.* (2002) *Mol. Cell* 10, 995–1005.
160. Sloan, K. E. *et al.* (2013) *Cell Rep.* 5, 237–247.
161. Brown, L. *et al.* (2007) *Crit. Rev. Eukaryot. Gene Expr.* 17, 73–85.
162. Jin, M. (2019) *Exp. Mol. Med.* 51, 1–10.
163. Kang, R. & Tang, D. (2017) *Curr. Pathobiol. Rep.* 5, 153–159.
164. Lee, J.-A. & Gao, F.-B. (2008) *Autophagy* 4, 230–232.
165. Zhao, M. *et al.* (2020) *Front. Med.* 7, 591736.
166. Tomar, D. *et al.* (2012) *Biochim. Biophys. Acta* 1823, 316–326.
167. Fujisawa, K. *et al.* (2016) *J. Exp. Clin. Cancer Res. CR* 35, 48.
168. Wolff, N. C. *et al.* (2011) *Mol. Cell. Biol.* 31, 1870–1884.
169. Kay, H. H. *et al.* (2007) *Placenta* 28, 854–860.
170. Kuo, M.-L. *et al.* (2016) *Sci. Rep.* 6, 18846.
171. Simon, N. E. *et al.* (2017) *Cell Cycle Georget. Tex* 16, 1175–1180.
172. Huang, N. *et al.* (2020) *Cell Death Dis.* 11, 507.
173. Woodman, P. (2016) *Biochem. J.* 473, e5-8.
174. Baghdadi, M. *et al.* (2018) *J. Leukoc. Biol.* 104, 931–951.
175. Xie, Z. *et al.* (2016) *AAPS J.* 18, 294–304.
176. Huang, B. & Baek, S.-H. (2017) *Mol. Pharmacol.* 91, 307–316.
177. Takei, Y. *et al.* (2002) *J. Biol. Chem.* 277, 43121–43125.
178. Vietri, M. *et al.* (2015) *Nature* 522, 231–235.
179. Niwa, S. (2015) *Anat. Sci. Int.* 90, 1–6.
180. Samora, C. P. *et al.* (2011) *Nat. Cell Biol.* 13, 1040–1050.
181. Martino, L. *et al.* (2017) *Dev. Cell* 43, 157-171.e7.
182. Hartono, null *et al.* (2019) *Cell Div.* 14, 7.
183. Pinto-Costa, R. & Sousa, M. M. (2020) *Cytoskelet. Hoboken NJ* 77, 76–83.
184. Shiratsuchi, G. *et al.* (2015) *EMBO J.* 34, 97–114.
185. Gabrielson, M. *et al.* (2016) *Biochem. Biophys. Res. Commun.* 469, 1090–1096.
186. Schulze, J. O. *et al.* (2008) *FEBS J.* 275, 4627–4640.
187. Maeda, M. *et al.* (2011) *Mol. Biol. Cell* 22, 3840–3852.
188. Peng, J.-M. *et al.* (2018) *Hepatol. Baltim. Md* 67, 2226–2243.
189. Florindo, C. *et al.* (2012) *J. Cell Sci.* 125, 3085–3090.
190. Noma, T. (2005) *J. Med. Investig. JMI* 52, 127–136.
191. Zhao, L. N. *et al.* (2021) *Oncogene* 40, 2339–2354.
192. Pedley, A. M. & Benkovic, S. J. (2017) *Trends Biochem. Sci.* 42, 141–154.
193. Kumari, N. *et al.* (2019) *Proc. Natl. Acad. Sci. U. S. A.* 116, 22140–22151.
194. Conaway, R. C. *et al.* (2005) *Trends Biochem. Sci.* 30, 250–255.
195. Finn, R. M. *et al.* (2012) *FASEB J.* 26, 4788–4804.
196. Pascual, M. *et al.* (2006) *Differ. Res. Biol. Divers.* 74, 65–80.
197. Miao, X. *et al.* (2020) *Biol. Reprod.* 103, 13–23.
198. Zhao, T. *et al.* (2012) *Cell Cycle Georget. Tex* 11, 4517–4523.
199. Han, H. *et al.* (2013) *Nature* 498, 241–245.
200. Motoi, N. *et al.* (2008) *Dev. Growth Differ.* 50, 307–320.
201. Hwang, I.-Y. *et al.* (2016) *Cell Metab.* 24, 494–501.
202. Zhao, P. *et al.* (2020) *Epigenetics* 15, 369–385.
203. Chen, G. *et al.* (2018) *Biochem. Biophys. Res. Commun.* 501, 259–265.
204. Tan, B. S. N. *et al.* (2016) *Mech. Dev.* 141, 32–39.
205. Kaehler, C. *et al.* (2012) *PloS One* 7, e50134.
206. White, E. J. F. *et al.* (2017) *Wiley Interdiscip. Rev. RNA* 8,.
207. Bardoni, B. *et al.* (2003) *Hum. Mol. Genet.* 12, 1689–1698.
208. Guo, M. & Schimmel, P. (2013) *Nat. Chem. Biol.* 9, 145–153.
209. Sankaranarayanan, R. & Moras, D. (2001) *Acta Biochim. Pol.* 48, 323–335.
210. Combs, D. J. *et al.* (2006) *Mol. Cell. Biol.* 26, 523–534.
211. Huang, N. *et al.* (2005) *J. Biol. Chem.* 280, 5496–5502.
212. O'Leary, M. N. *et al.* (2013) *PLoS Genet.* 9, e1003708.
213. Kuwata, H. & Hara, S. (2019) *Prostaglandins Other Lipid Mediat.* 144, 106363.
214. Pegg, A. E. (2009) *IUBMB Life* 61, 880–894.
215. Prabhudesai, S. *et al.* (2018) *Front. Cell Dev. Biol.* 6, 14.
216. Vockley, J. & Ensenauer, R. (2006) *Am. J. Med. Genet. C Semin. Med. Genet.* 142C, 95–103.
217. Swanson, D. A. *et al.* (2001) *Mol. Cell. Biol.* 21, 1058–1065.
218. El-Hattab, A. W. (2016) *Mol. Genet. Metab.* 118, 153–159.
219. Adeva-Andany, M. M. *et al.* (2016) *BBA Clin.* 5, 85–100.
220. Wichelecki, D. J. *et al.* (2014) *Biochemistry* 53, 2732–2738.
221. Aydemir, T. B. *et al.* (2016) *J. Biol. Chem.* 291, 23939–23951.
222. Arif, A. *et al.* (2017) *Nature* 542, 357–361.
223. van den Boomen, D. J. H. *et al.* (2020) *Curr. Opin. Cell Biol.* 65, 103–111.
224. Lanning, N. J. *et al.* (2014) *Cell Rep.* 7, 907–917.
225. Gakh, O. *et al.* (2002) *Biochim. Biophys. Acta* 1592, 63–77.
226. Safaee, M. *et al.* (2013) *Int. J. Oncol.* 43, 1343–1350.
227. Boersma, H. H. *et al.* (2005) *J. Nucl. Med. Off. Publ. Soc. Nucl. Med.* 46, 2035–2050.
228. Giménez-Mascarell, P. *et al.* (2018) *Front. Pediatr.* 6, 77.
229. Tomsig, J. L. *et al.* (2003) *J. Biol. Chem.* 278, 10048–10054.
230. Kondratiuk, I. *et al.* (2020) *Proc. Natl. Acad. Sci. U. S. A.* 117, 13468–13479.
231. Johnson, E. N. & Druey, K. M. (2002) *J. Biol. Chem.* 277, 16768–16774.
232. Bouter, A. *et al.* (2015) *Placenta* 36 Suppl 1, S43-49.
233. Hanna, M. G. *et al.* (2018) *BioEssays News Rev. Mol. Cell. Dev. Biol.* 40, e1800004.
234. Lerner, M. *et al.* (2007) *Mol. Biol. Cell* 18, 1670–1682.
235. Myllyharju, J. & Kivirikko, K. I. (2004) *Trends Genet. TIG* 20, 33–43.
236. Kay, E. J. *et al.* (2021) *Front. Oncol.* 11, 719922.
237. Schrankel, C. S. & Hamdoun, A. (2021) *Dev. Biol.* 472, 115–124.
238. Beck, M. & Hurt, E. (2017) *Nat. Rev. Mol. Cell Biol.* 18, 73–89.
239. Smith, D. E. *et al.* (2013) *Mol. Aspects Med.* 34, 323–336.
240. Hetzer, M. *et al.* (2002) *Nat. Cell Biol.* 4, E177-184.
241. Chook, Y. M. & Süel, K. E. (2011) *Biochim. Biophys. Acta* 1813, 1593–1606.
242. Taylor, K. M. *et al.* (2005) *FEBS Lett.* 579, 427–432.
243. Mandò, C. *et al.* (2013) *Pediatr. Res.* 74, 104–110.
244. Xu, X. *et al.* (2020) *J. Bone Miner. Res. Off. J. Am. Soc. Bone Miner. Res.* 35, 116–129.
245. Tomsig, J. L. & Creutz, C. E. (2002) *Cell. Mol. Life Sci. CMLS* 59, 1467–1477.
246. Bill, C. A. & Vines, C. M. (2020) *Adv. Exp. Med. Biol.* 1131, 215–242.
247. Chen, J.-J. *et al.* (2019) *Theranostics* 9, 5049–5064.
248. Jiang, H. *et al.* (2020) *BioMed Res. Int.* 2020, 3418398.
249. Duan, S. *et al.* (2018) *J. Exp. Clin. Cancer Res. CR* 37, 304.
250. Totaro, A. *et al.* (2018) *Nat. Cell Biol.* 20, 888–899.
251. Wu, X. *et al.* (2020) *Biochem. Biophys. Res. Commun.* 527, 29–36.
252. Momiuchi, Y. *et al.* (2015) *J. Biol. Chem.* 290, 23816–23825.
253. Tang, H. *et al.* (2021) *Front. Genet.* 12, 689097.
254. Hansson, S. R. *et al.* (2006) *Mol. Hum. Reprod.* 12, 169–179.
255. Shen, H. *et al.* (2020) *Reprod. Sci. Thousand Oaks Calif* 27, 1553–1561.
256. Schumann, S. *et al.* (2015) *Histochem. Cell Biol.* 144, 571–585.
257. Domino, S. E. *et al.* (2001) *Mol. Cell. Biol.* 21, 8336–8345.
258. Highet, A. R. *et al.* (2015) *Eur. J. Cell Biol.* 94, 589–602.
259. Lindau, R. *et al.* (2018) *Hum. Reprod. Oxf. Engl.* 33, 588–599.
260. Shalom-Barak, T. *et al.* (2012) *Dev. Biol.* 372, 143–155.
261. Xiao, L.-J. *et al.* (2006) *Reprod. Camb. Engl.* 131, 1137–1149.
262. J, S. *et al.* (2019) *Am. J. Transl. Res.* 11,.
263. Rokavec, M. *et al.* (2017) *Sci. Rep.* 7, 4687.
264. Alcantara, O. *et al.* (1994) *Biochem. J.* 298 ( Pt 2), 421–425.
265. Zhao, N. *et al.* (2010) *J. Biol. Chem.* 285, 32141–32150.
266. Tzima, E. & Schimmel, P. (2006) *Trends Biochem. Sci.* 31, 7–10.
